# Sirolimus plus nintedanib treats vascular pathology in HHT mouse models

**DOI:** 10.1101/739144

**Authors:** Santiago Ruiz, Haitian Zhao, Pallavi Chandakkar, Julien Papoin, Hyunwoo Choi, Aya Nomura-Kitabayashi, Radhika Patel, Matthew Gillen, Li Diao, Prodyot K. Chatterjee, Mingzhu He, Yousef Al-Abed, Ping Wang, Christine N. Metz, S. Paul Oh, Lionel Blanc, Fabien Campagne, Philippe Marambaud

## Abstract

Hereditary hemorrhagic telangiectasia (HHT), a genetic bleeding disorder leading to systemic arteriovenous malformations (AVMs), is caused by loss-of-function mutations in the ALK1-ENG-Smad1/5/8 pathway. Evidence suggests that HHT pathogenesis strongly relies on overactivated PI3K-Akt-mTOR and VEGFR2 pathways in endothelial cells (ECs). In the BMP9/10-immunoblocked (BMP9/10ib) neonatal mouse model of HHT, we report here that the mTOR inhibitor, sirolimus, and the receptor tyrosine-kinase inhibitor, nintedanib, could synergistically fully block, but also reversed, retinal AVMs to avert retinal bleeding and anemia. Sirolimus plus nintedanib prevented vascular pathology in the oral mucosa, lungs, and liver of the BMP9/10ib mice, as well as significantly reduced gastrointestinal bleeding and anemia in inducible ALK1-deficient adult mice. Mechanistically, in vivo in BMP9/10ib mouse ECs, sirolimus and nintedanib blocked the overactivation of mTOR and VEGFR2, respectively. Furthermore, we found that sirolimus activated ALK2-mediated Smad1/5/8 signaling in primary ECs—including in HHT patient blood outgrowth ECs—and partially rescued Smad1/5/8 activity in vivo in BMP9/10ib mouse ECs. These data demonstrate that the combined correction of endothelial Smad1/5/8, mTOR, and VEGFR2 pathways opposes HHT pathogenesis. Repurposing of sirolimus plus nintedanib might provide therapeutic benefit in HHT patients.

## INTRODUCTION

Hereditary hemorrhagic telangiectasia (HHT), also known as Rendu-Osler-Weber syndrome, is an autosomal dominant genetic disease with a prevalence of ∼1 in 5,000 individuals (1). HHT is a hemorrhagic vascular dysplasia characterized by the development of mucocutaneous telangiectasias and arteriovenous malformations (AVMs) in multiple tissues and organs. Particularly affected is the vasculature of the liver, lungs, and oronasal and gastrointestinal (GI) mucosa. In its most severe manifestations, HHT can lead to highly debilitating and life-threatening events, such as severe epistaxis and internal bleeding. HHT is also associated with secondary complications, which include anemia, cerebral abscess and embolism following pulmonary AVMs, as well as high-output cardiac failure consecutive to liver AVMs (2, 3).

At least 85% of HHT patients carry disease-causing loss-of-function mutations in the genes, *ENG* (encoding endoglin) or *ACVRL1* (encoding activin receptor-like kinase 1, ALK1), which define the two disease subtypes: HHT1 (OMIM #187300) and HHT2 (#600376), respectively (4, 5). Mutations were also reported in *SMAD4* (6) (encoding Smad4) and *GDF2* (7) (encoding bone morphogenetic protein 9, BMP9), which cause rare and atypical forms of the disease called juvenile polyposis/HHT combined syndrome (OMIM #175050) and HHT-like vascular anomaly syndrome (#615506), respectively. BMP9, ALK1, endoglin, and Smad4 are members of the transforming growth factor-β signaling superfamily and all functionally interact in the same signal transduction axis (8). The cell surface receptor complex composed of the co-receptor endoglin, the endothelial BMP type I receptor ALK1, and a BMP type II receptor (e.g., BMPR2) is activated by sequential binding to the circulating ligands BMP9 and BMP10 (9–11). Mutations in *BMPR2* cause familial pulmonary arterial hypertension (PAH), which can be observed in some HHT2 patients (12), further supporting the notion that ALK1 and BMPR2 functionally interact. ALK1-endoglin receptor activation leads to phosphorylation of the signal transducers, Smad1, Smad5, and Smad8, to trigger the formation of Smad1/5/8-Smad4 complexes that translocate into the nucleus to control specific gene expression programs (13).

HHT pathogenesis is triggered by a reduction of Smad1/5/8 signaling in ECs. Indeed, HHT-causing mutations decrease Smad1/5/8 response to ALK1-endoglin receptor activation by BMP9 (14–16). Consistent with this model, ALK1, endoglin, or Smad4 inactivation in mice and zebrafish leads to vascular defects, which include hypervascularization and AVMs (17–25). Downstream from ALK1-endoglin receptor loss-of-function, the exact pathways involved in HHT pathogenesis and ultimately, AVM development—i.e., the formation of direct shunts between an artery and a vein—remain incompletely understood (26). However, concordant studies have shown that HHT is associated with abnormal reactivation of angiogenesis (27) and that overactivated proangiogenic pathways, such as VEGF signaling, are required for the development of the vascular pathology of HHT models (28, 29). In vitro data in cell cultures have further revealed that VEGFR2 phosphorylation and activity were increased upon ALK1 silencing (30), while endoglin silencing affected VEGFR2 trafficking to increase its downstream signaling (31). Transcriptomic data in ECs have also demonstrated that ALK1 negatively controlled VEGFR2 (*KDR* gene) expression (32, 33). In vivo, *Vegfr2* knockdown was reported to block hypervascularization and AVMs in *Alk1*^iΔEC^ mice (30) and pharmacological inhibition of VEGFR2 reduced disease severity in *Eng*^iΔEC^ mice (31). Also, recent clinical trials have indicated that intravenous treatment with the VEGF blocking antibody, bevacizumab, provided some therapeutic benefits in severely affected HHT patients (34, 35). In summary, although the field has not reached a consensus on the exact mechanism by which ALK1-endoglin loss-of-function activates VEGF signaling, there is strong evidence that VEGF-VEGFR2 signaling is a key contributor of the pathogenic process of HHT.

PI3K-Akt overactivation is also involved in AVM development in ALK1 (30) and endoglin (31) deficient mice and in HHT2 patients (36). Specifically, ribosomal protein S6 (S6) phosphorylation, a downstream effector of mTORC1 and Akt, was reported to be increased in the ECs of AVMs in the *Alk1*^iΔEC^ mouse retina and in BMP9- and BMP10-immunoblocked (BMP9/10ib) mice (30), another HHT model. In addition, administration of PI3K inhibitors, such as wortmannin, partially reduced retinal AVM numbers in *Alk1*^iΔEC^, *Eng*^iΔEC^, and *Smad4*^iΔEC^ mice (25, 30, 31). Akt1 deficiency also mitigated AVM pathology in the *Smad4*^iΔEC^ model (25). Thus, overactivated VEGFR2 and PI3K-Akt-mTOR pathways are critical drivers—in addition to ALK1-endoglin loss-of-function—of HHT pathogenesis and AVM development.

Recently, we (32) and others (37) have reported that tacrolimus (Tac) was associated with Smad1/5/8 signaling activating properties, both in vitro in primary ECs and in vivo in mouse models of HHT and PAH. Spiekerkoetter *et al.* further reported in a cell-based BMP/Smad transactivation assay and in human pulmonary arterial ECs, that the mTOR inhibitor sirolimus (Siro, rapamycin), a macrolide analog of Tac, also activated Smad1/5/8, although less potently than Tac (37). These data suggest the intriguing possibility that Siro could both prevent mTOR overactivation and rescue Smad1/5/8 signaling in HHT models. In line with this hypothesis, a 2006 case study reported the potential beneficial effect of an immunosuppressive regimen, which contained Siro, in a probable HHT patient who underwent liver transplantation following high-output cardiac failure and hepatic AVM development (38). In this patient, the authors observed a reduction in angiodysplasia and mucosal hemorrhage. They proposed that the combination of Siro plus aspirin, in the antirejection regimen, could have contributed to the observed effects in this HHT patient (38).

Several receptor tyrosine kinase (RTK) inhibitors that target VEGFR2 are FDA/EMA-approved, including pazopanib that showed beneficial effects in ALK1 deficient mice (39) and in a cohort of HHT patients (40). A recent case study reported that nintedanib (Nin, BIBF 1120), another VEGFR2-targeting RTK inhibitor, which is approved for idiopathic pulmonary fibrosis (IPF) treatment, produced marked reductions in epistaxis and skin telangiectasias in a patient affected by both HHT and IPF (41). Nin is of particular interest compared to other classical RTK inhibitors because it has a narrow spectrum of inhibition and is associated with minimal adverse effects.

In this context and in the current study, we sought to determine whether concurrent pharmacological targeting of Smad1/5/8 reduction, and of mTOR and VEGFR2 activation, could rescue the signaling defects and vascular pathology of HHT. We report that Siro and Nin, when administered in combination, mechanistically synergize at the transcriptional and signaling levels to efficiently prevent and reverse the vascular pathology, and the associated bleeding and anemia, in two HHT mouse models.

## RESULTS

### Sirolimus and nintedanib combination prevents vein dilation, hypervascularization, and AVM development in the retina of the tBMP9/10ib mice

Recently, we have reported the development of a HHT mouse model, which uses the transmammary route for administering BMP9 and BMP10 blocking antibodies to nursing mouse pups (22, 32), hereafter referred to as the transmammary-treated BMP9/10 immunoblocked (tBMP9/10ib) mice. We showed that this model, which has the advantage of being less invasive for the pups and more suitable for the analysis of large cohorts, leads to reduced ALK1 signaling and the development of a robust HHT-like pathology in the retinal vasculature of the pups (22, 32). In this model, we found that Tac significantly activated endothelial ALK1 signaling in vivo and prevented hypervascularization of the tBMP9/10ib retina. Further analyses of Tac efficacy in preventing vascular pathology in tBMP9/10ib mice revealed, however, a relatively modest effect of the drug on AVM development (32). These data prompted us to determine whether the macrolide analog of Tac, Siro (administered alone and in combination with Nin), is more potent in reducing the vascular pathology of the tBMP9/10ib retina.

Studies were conducted prior to these experiments to determine the appropriate dosing and injection schedule of the drugs in pups. The highest dose of Siro that did not affect normal vascular development was chosen [0.5 mg/kg, intraperitoneal (i.p.) injection of the pups, assessed in the neonatal retina]. Similarly, in order to prevent changes in retinal vascular development, we determined that Nin should be given every third day and not before P5. With this schedule for Nin injection, the highest dose of the drug that did not affect physiological vascular development was determined to be 0.3 mg/kg (i.p., data not shown). Co-administration of Siro and Nin (Siro+Nin) at these dosing and injection schedules did not also significantly affect normal vascular growth, indicating that physiological angiogenesis in the retina was not inhibited by the drug combination (Figure S1). At this dosing, LC-MS analyses measured average serum concentrations of 9.0 nM Siro and 5.3 nM Nin in the injected P6 pups.

As shown in Figure 1A and described before (22, 32), vascular pathology in pups was initiated at postnatal day 3 (P3) by one i.p. injection of the dams with anti-BMP9/10 antibodies. Mouse pups were administered preventively and daily with Siro from P3 to P5 (0.5 mg/kg, i.p.) and with Nin at P5 (0.3 mg/kg, i.p.). Mice were then analyzed at P6 (Figure 1A), a time point at which retinal vessel dilation, hypervascularization, and AVMs can readily be observed and quantified in this model (22, 32). In the tBMP9/10ib retinas, Siro significantly reduced AVM number and AVM diameter (Figures 1B-1D, 1G-1I, 1Q, and 1R). In addition, as we observed previously for Tac at the same dosing (0.5 mg/kg, P3-P5, i.p.) (32), Siro prevented vein dilation (Figure 1S) and the increase in density of the vascular plexus (Figures 1B-1D, 1L-1N, and 1T). In contrast, Nin at the tested dosing failed to reduce any of the investigated vascular defects of the tBMP9/10ib retinas (Figures 1B, 1C, 1E, 1G, 1H, 1J, 1L, 1M, 1O, and 1Q-1T).

**Figure 1.**
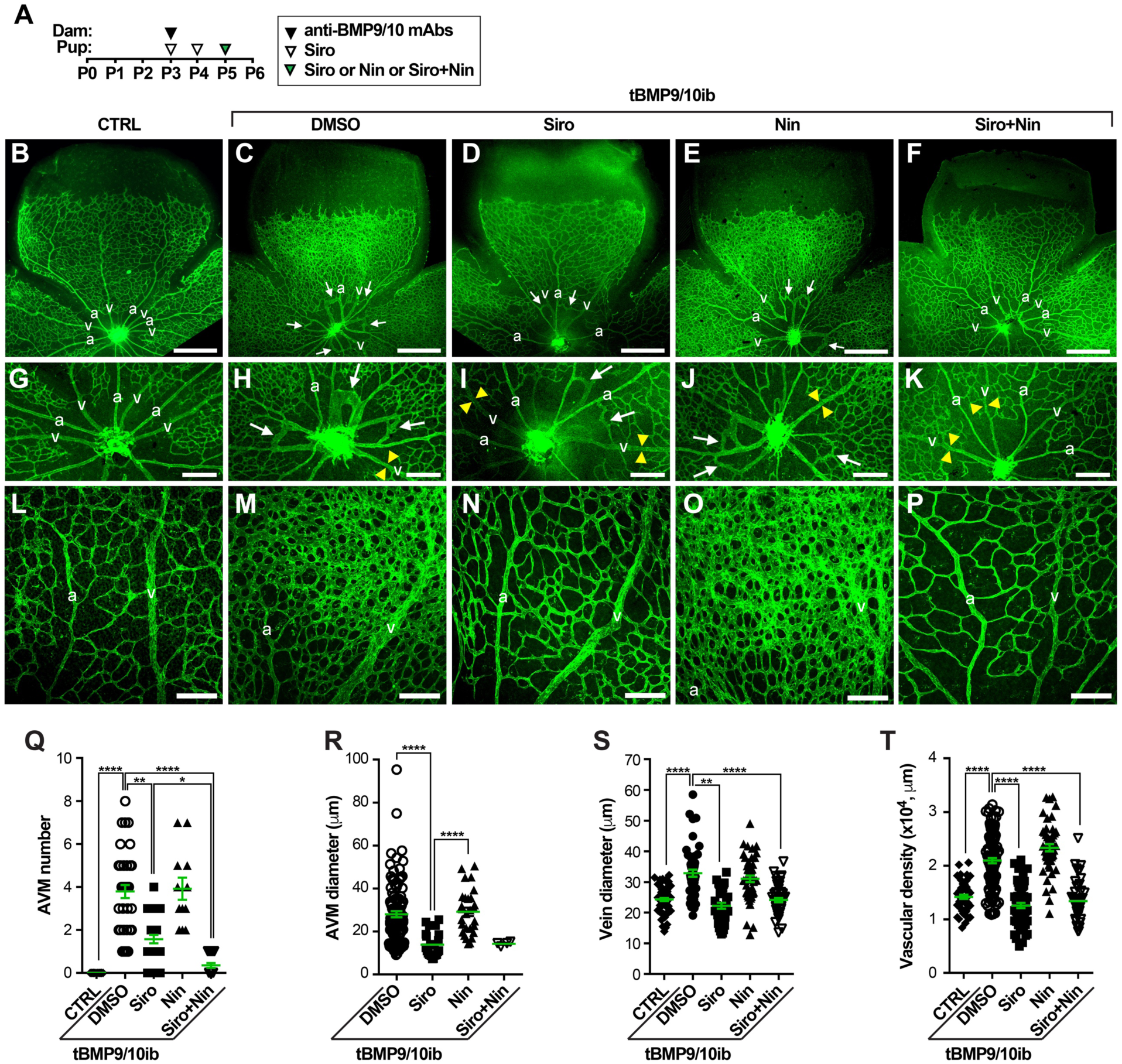
Siro+Nin prevents vein dilation, hypervascularization, and AVMs in the tBMP9/10ib mouse retina. **(A)** Schematic representation of the protocol of disease induction (i.p. injections of the lactating dams with anti-BMP9/10 blocking antibodies, anti-BMP9/10 mAbs) and of drug treatments (i.p. injections of the pups with Siro and/or Nin). Arrowheads indicate the postnatal days of injection. Control mouse pups were obtained by injecting the lactating dams with isotype controls and by directly injecting them with DMSO. Pups were euthanized at P6. **(B-P)** Representative images of retinas stained with fluorescent isolectin B4 (green) from control (CTRL; B, G, and L) mice and tBMP9/10ib mice treated with vehicle (DMSO; C, H, and M), Siro (D, I, and N), Nin (E, J, and O), or the combination of Siro and Nin (F, K, and P). Higher magnification in (G-K) and (L-P) show respectively retinal vein diameter (yellow arrowheads) and retinal vascular fields (plexus area) between an artery (a) and a vein (v). Arrows in denote AVMs. Scale bars, 500 µm (B-F), 200 µm (G-K), and 100 µm (L-P). **(Q-T)** Scatter plots measuring AVM number (Q), AVM diameter (R), vein diameter (S) and vascular plexus density (T) in CTRL and tBMP9/10ib mice treated as in (B-F). Data represent individual retinas and mean ± SEM (n = 14, 38, 28, 12, and 20 retinas for the CTRL, DMSO, Siro, Nin, and Siro+Nin groups, respectively); Kruskal-Wallis test, post hoc Dunn’s multiple comparison test; \**P*<0.05, \*\**P*<0.01, \*\*\*\**P*<0.0001.

Although Siro treatment was able to significantly reduce AVM number and size, its preventive effect was only partial (AVM number, mean = 3.79 ± 0.30 in DMSO-treated tBMP9/10ib retinas vs. mean = 1.57 ± 0.19 in Siro-treated tBMP9/10ib retinas, *P*≤0.01). Knowing that evidence is strong to suggest that VEGFR2 signaling is increased in HHT models and that Siro demonstrated no efficacy in inhibiting VEGFR2-mediated MAPK signaling activation in primary ECs [Figure S2 and Ref. (42)], we asked whether the VEGFR2 inhibitor Nin could increase Siro potency in preventing AVMs. Combination treatment with the two drugs resulted in a significant increase of Siro anti-AVM effect (Figures 1B-1K and 1Q; AVM number after treatment, mean = 0.35 ± 0.11, *P*≤0.0001 vs. DMSO-treated tBMP9/10ib retinas, and *P*≤0.05 vs. Siro-treated tBMP9/10ib retinas). Siro+Nin combination did not further increase the effect of Siro on vein dilation and vascular density, as Siro alone was sufficient to fully correct these two defects (Figures 1L-1P, 1S, and 1T). Measurement of the diameter of the few remaining AVMs identified in the retina of the Siro+Nin-treated mice also revealed no difference compared to treatment with Siro alone (Figure 1R). Together these data in tBMP9/10ib mice show that Siro fully normalized vein dilation and hypervascularization, and significantly lowered AVM number and size in the retina. Furthermore and more strikingly, Nin significantly strengthened the anti-AVM effect of Siro.

### Siro+Nin prevents anemia and retinal bleeding in tBMP9/10ib mice

We next assessed pathology progression in tBMP9/10ib pups at P9. As before, vascular pathology in pups was initiated at P3 by one i.p. injection of the dams with anti-BMP9/10 antibodies (Figure 2A). Complete blood count (CBC) revealed significant reductions in hematocrit level, red blood cell (RBC) number, and hemoglobin level (Figure 2B), indicative of anemia in P9 tBMP9/10ib pups. Furthermore, severe cardiomegaly (Figures 2C and 2D) and splenomegaly (Figure 2C) developed in tBMP9/10ib mice. Splenomegaly was accompanied by an expansion of the red pulp (Figure 2D, right panels), indicating the presence of splenic erythropoietic stress response consecutive to anemia. These data prompted us to investigate whether tBMP9/10ib mice were actively bleeding. Inspection of the retinas using whole-mount immunohistochemistry (IHC) with an antibody directed against the RBC marker, Ter119, revealed the presence of strongly immunoreactive patches in multiple areas of the tBMP9/10ib retinas (Figures 2E, 2F, and 2K). Single-cell resolution confocal analyses and 3D reconstruction showed the presence of RBC patches outside the tBMP9/10ib retinal vasculature (Figures 2H-2H” and 2I-2I”, and animation in Movie S1). Interestingly, isolectin B4-positive projections could clearly be identified near the center of some of these RBC accumulations, suggesting that they might represent transversal vascular projections at the origin of the bleeding (Figures 2I-2I”, arrows; Movie S1). Treatment of the tBMP9/10ib mice with Siro+Nin from P3 to P8 (Figure 2A) significantly reduced the area occupied by retinal bleeding (Figures 2E-2K) and fully prevented the decrease in hematocrit level, RBC number, and hemoglobin level (Figure 2B), as well as blocked cardiomegaly, splenomegaly, and the loss of splenic architecture (Figures 2C and 2D). Thus, Siro+Nin combination treatment prevented anemia and retinal bleeding in tBMP9/10ib mice.

**Figure 2.**
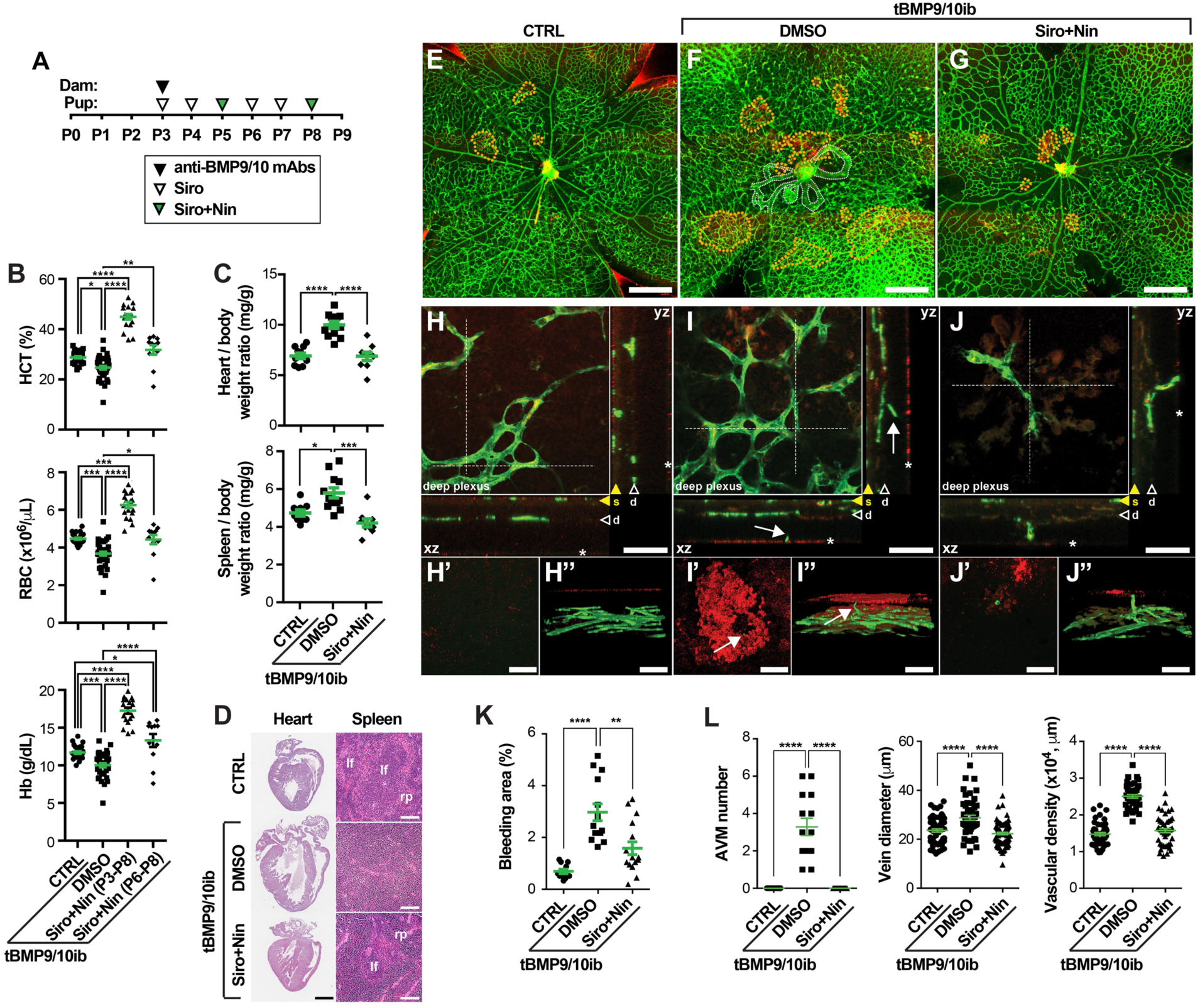
Siro+Nin prevents anemia and retinal bleeding in tBMP9/10ib mice. **(A)** Schematic representation of the protocol of disease induction and drug treatments. Pups were euthanized at P9. **(B)** Scatter plots measuring hematocrit (HCT) level, red blood cell (RBC) number, and hemoglobin (Hb) level in P9 control (CTRL) and tBMP9/10ib mice treated with Siro+Nin at P3-P8 [as in (A)] or P6-P8 (see Figure 3A). Data represent individual mice and mean ± SEM (n = 26-29, 35-40, 18, 11 mice for the CTRL, DMSO, Siro+Nin (P3-P8), and Siro+Nin (P6-P8) groups, respectively); HCT and RBC analyses: Kruskal-Wallis test, post hoc Dunn’s multiple comparison test; Hb analysis: one-way ANOVA, Tukey’s multiple comparison test. **(C and D)** P9 mouse heart and spleen analyses of CTRL and tBMP9/10ib mice treated as in (A). Heart/body and spleen/body weight ratios are shown in (C). Data represent individual mice and mean ± SEM (n = 9-11, 12, 9 mice for the CTRL, DMSO, and Siro+Nin groups, respectively); one-way ANOVA, Tukey’s multiple comparison test. Heart and spleen sections stained with H&E are shown in (D). lf, lymphoid follicle; rp, red pulp. **(E-J”)** Representative images showing microbleeds (dotted orange lines in E-G) in retinas stained with fluorescent isolectin B4 (green) and anti-Ter119 (red) from CTRL (E, H-H”), and tBMP9/10ib mice treated with DMSO (F, I-I”) or Siro+Nin (G, J-J”), as in (A). Dotted white lines in (F) indicate AVMs. Higher magnifications in (H-J”) show the extending deeper retinal plexus [arrowheads marked as “d” in (H, I, and J)] and orthogonal view (xz and yz planes) of the stack of images at the level of the dotted lines shown in (H, I, and J). Orthogonal views in (H, I, and J) show superficial vascular plexus (yellow arrowheads marked as “s”), vascular branches projecting to the outer layer of the retina (white arrows), and microbleeds (asterisks). Focal plane views (H’, I’, and J’) and 3D reconstructions (H”, I”, and J”) identify microbleeds. Scale bars, 1 mm (D, Heart), 10 µm (D, Spleen), 500 µm (E-G), and 50 µm (H-J”). **(K)** Scatter plot measuring bleeding areas, expressed as a percentage of the retinal vascular area occupied by extravascular accumulation of RBCs in mice treated as in (A). Data represent mean ± SEM (n = 6, 4, 4 mice for the CTRL, DMSO, and Siro+Nin groups, respectively); one-way ANOVA, Tukey’s multiple comparison test. **(L)** Scatter plots measuring AVM number, vein diameter, and vascular plexus density in the retina of P9 pups treated as in (A). Data represent individual retinas and mean ± SEM (n = 7, 7, 8 mice for the CTRL, DMSO, and Siro+Nin groups, respectively); AVM number and vein diameter analyses: one-way ANOVA, Tukey’s multiple comparison test; vascular density analysis: Kruskal-Wallis test, post hoc Dunn’s multiple comparison test. \**P*<0.05, \*\**P*<0.01, \*\*\**P*<0.001, \*\*\*\**P*<0.0001.

Because evidence is mounting that AVM development is controlled by blood flow (43) and because P9 tBMP9/10ib mice display significant cardiomegaly, we asked: (i) whether tBMP9/10ib mice develop cardiac defects leading to higher cardiac output and (ii) whether the drugs could indirectly reduce vascular pathology by correcting these cardiac defects. Although a modest decrease in heart rate was measured by doppler ultrasonography upon drug treatment in tBMP9/10ib mice, heart rate was overall not significantly changed in DMSO-treated and Siro+Nin-treated tBMP9/10ib pups, compared to normal pups (Figure S3A). In addition, no significant defects in cardiac output (Figure S3B) or pulmonary arterial pressure (Figure S3C) measures were found between all groups. These data demonstrate that basic cardiac function is normal in tBMP9/10ib pups, at least until P9, and that Siro+Nin combination is therefore unlikely to act on the vasculature by changing cardiac output.

### Siro+Nin reverses vascular pathology in tBMP9/10ib mice

We next analyzed the retinal vasculature of P9 tBMP9/10ib mice treated or not with Siro+Nin. On average ∼4 AVMs were detected in tBMP9/10ib mice (Figures 2L and S4), indicating that no additional AVMs developed between P6 (mean = 3.79 ± 0.30, Figure 1Q) and P9. Strikingly, while Siro+Nin-treated P6 mice still contained some AVMs (n = 0.35 ± 0.11, Figure 1Q), the P9 tBMP9/10ib mouse retinas that were treated with the drugs for 3 additional days were devoid of AVMs (Figures 2L and S4). In addition, the 6-day-Siro+Nin treatment (P3-P8, Figure 2A), as we observed after the 3-day-Siro+Nin treatment (P3-P5, Figure 1), fully prevented vein dilation and hypervascularization (Figures 2L and S4). LC-MS analyses measured an increase in average drug concentrations in the pup serum between P6 and P9: from 9.0 nM to 22.6 nM Siro and from 5.3 nM to 24.7 nM Nin, indicating that three additional days of drug treatment led to an accumulation of the drugs in the circulation.

These data suggest that Siro+Nin treatment might also reverse existing AVMs, since some AVMs disappeared between P6 and P9. To directly address this possibility, we implemented a protocol that started the drug treatment once retinal vascular pathology was established. Specifically, pathology was induced as before at P3 and pups were then treated at P6 with Siro+Nin, a time point where we have established the presence of robust vein dilation, hypervascularization, and of ∼4 AVMs per retina (Figure 1). Pups were treated for 3 days, from P6 to P8 (P6-P8), and analyzed at P9 (Figure 3A). We found that Siro+Nin administered after pathology induction significantly reduced overall vascular pathology (Figure S5), including AVM number, AVM size, vein dilation, and vascular density (Figures 3B and 3C). In addition, P6-P8 Siro+Nin treatment significantly increased hematocrit level, RBC number, and hemoglobin level in anemic tBMP9/10ib mice (Figure 2B).

**Figure 3.**
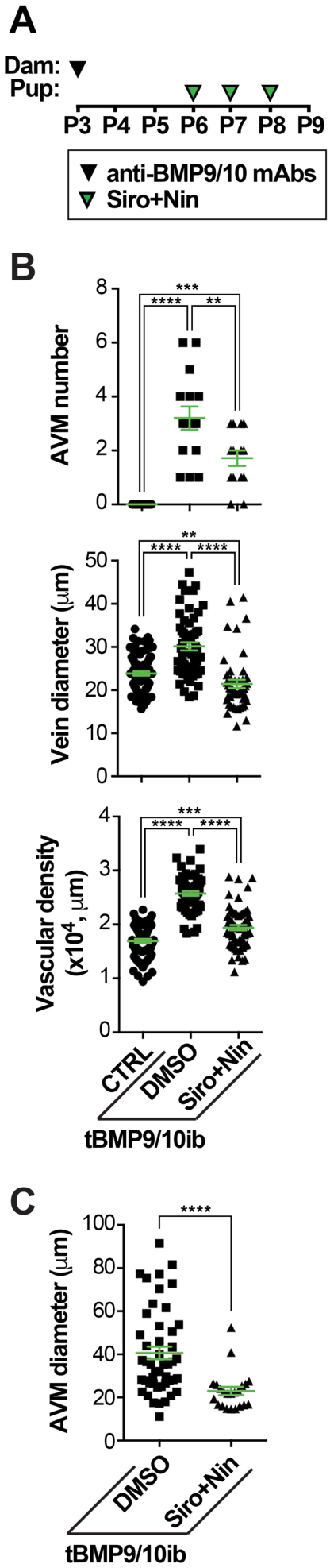
Siro+Nin reverses retinal vascular pathology in tBMP9/10ib mice. **(A)** Schematic representation of the protocol of disease induction and drug treatments. Pups were euthanized at P9. **(B and C)** Scatter plots measuring AVM number, vein diameter, vascular plexus density (B), and AVM diameter (C) in the retina of CTRL and tBMP9/10ib mice treated as in (A). Data represent individual retinas and mean ± SEM (n = 9, 8, 7 mice for the CTRL, DMSO, and Siro+Nin groups, respectively); AVM number and vascular density analyses: one-way ANOVA, Tukey’s multiple comparison test; AVM diameter analysis: Mann Whitney test; vein diameter analysis: Kruskal-Wallis test, post hoc Dunn’s multiple comparison test; \*\**P*<0.01, \*\*\**P*<0.001, \*\*\*\**P*<0.0001.

Since disease induction is triggered by only one i.p. injection at P3 of anti-BMP9/10 antibodies, we verified that the observed effects of Siro+Nin in P9 pups were not facilitated by a disappearance of the disease-causing anti-BMP9/10 blocking antibodies from the pup circulation. Using specific anti-BMP9 and anti-BMP10 antibody ELISAs, we found that serum antibody concentrations were stable between P6 and P9, and reached ∼70 µg/mL for the anti-BMP9 antibody and ∼85 µg/mL for the anti-BMP10 antibody (Figure S6). Thus, the disease-causing effects of the anti-BMP9/10 antibodies was maintained between P6 and P9, and therefore, the drug combination effect on pre-existing AVMs occurred in a maintained pathogenic environment. Taken together, these findings demonstrate that Siro+Nin combination treatment not only prevented, but also reversed, the retinal vascular pathology of the tBMP9/10ib mice.

### Siro+Nin prevents vascular pathology in the oral mucosa and lungs of the tBMP9/10ib mice

The oral mucosa and lungs are major sites of vascular lesion development in HHT patients. We investigated whether vascular defects are observed in these tissues of the P9 tBMP9/10ib mice. Injections of latex blue dye in the blood circulation were used to visualize vascular pathology. In the tBMP9/10ib mouse tongue and palate, mucosal vein dilation and hyperproliferative vascular defects were clearly identified after latex dye injection (Figures 4A, 4B, 4D, and 4E). Significant enlargements of the lingual and greater palatine vessels could be measured, compared to control tongues and palates (Figure 4J). In the lungs, the dye invaded a hypervascularized network of dilated small vessels throughout the lobar system and revealed an enlargement of the main pulmonary vessels of the tBMP9/10ib mice (Figures 4G, 4H, and 4J). Siro+Nin treatment of the tBMP9/10ib mice significantly and efficiently prevented the hyperproliferative vascular pathology and vessel dilation phenotype observed in the tongue, palate mucosa, and lungs (Figure 4). Thus, Siro+Nin combination reduced vascular pathology in the oral mucosa and lungs.

**Figure 4.**
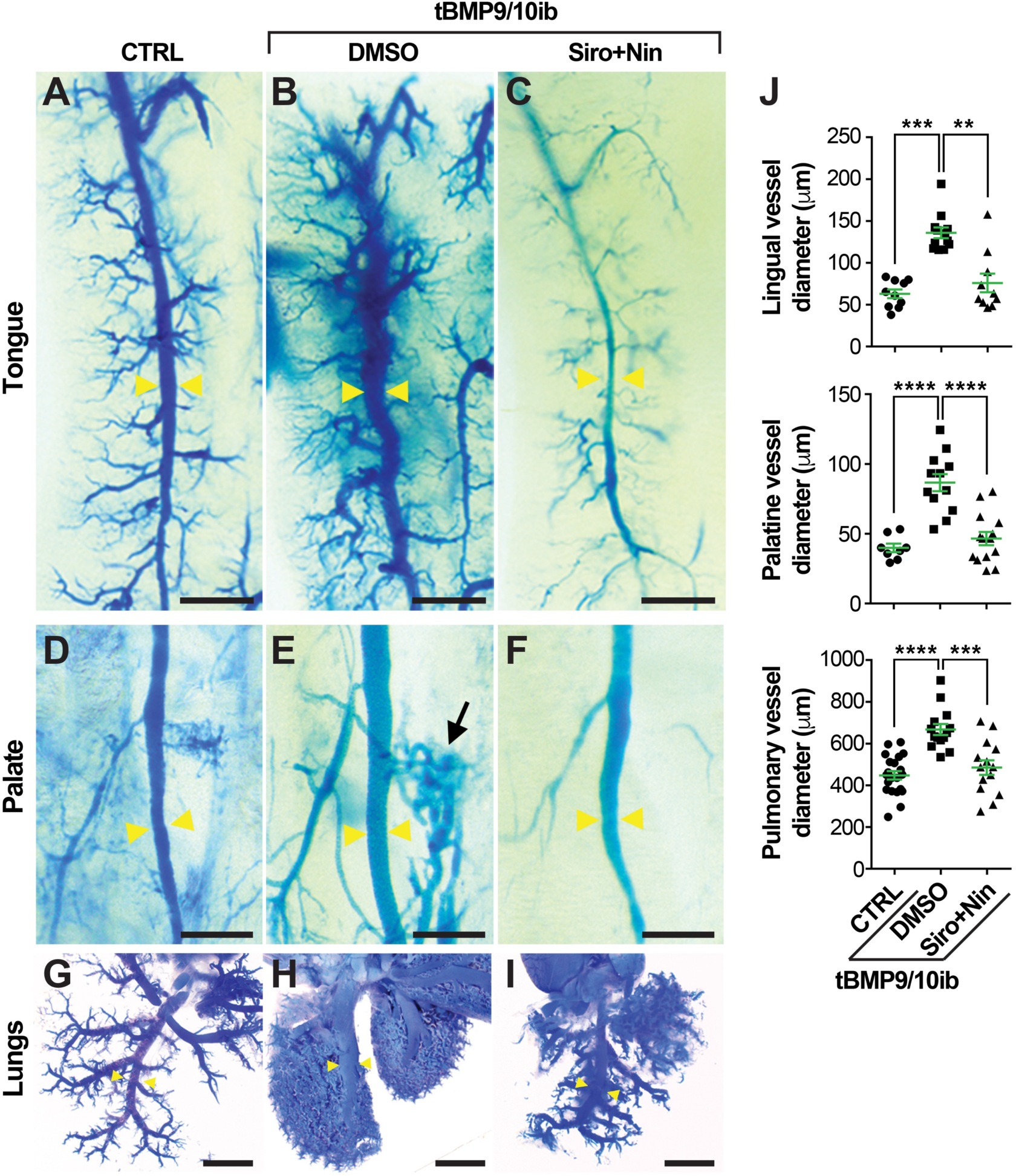
Siro+Nin prevents vascular pathology in the oral mucosa and lungs of the tBMP9/10ib mice. **(A-I)** Representative images of latex blue dye-perfused mouse tongue, palate, and lungs of control (CTRL; A, D, and G) and tBMP9/10ib mice treated with DMSO (B, E, and H) or Siro+Nin (C, F, and I), as in Figure 2A. Yellow arrowheads indicate lingual, palatine, and pulmonary vessels used for diameter measurements, black arrow denotes a dilation of a lateral branch in the palatine vasculature (E). Scale bars, 500 µm (A-F) and 2 mm (G-I). **(J)** Scatter plots measuring lingual, palatine, and pulmonary vessel diameter. Data represent mean ± SEM; lingual vessel diameter analysis (n = 3, 4, 3 mice for the CTRL, DMSO, and Siro+Nin groups, respectively): Kruskal-Wallis test, post hoc Dunn’s multiple comparison test; Palatine (n = 2, 3, 4 mice) and pulmonary (n = 10, 6, 6 mice) vessel diameter analyses: one-way ANOVA, Tukey’s multiple comparison test; \*\**P*<0.01, \*\*\**P*<0.001, \*\*\*\**P*<0.0001.

### Siro+Nin corrects a gene expression signature and prevents vascular pathology in the liver of the tBMP9/10ib mice

The liver is the most vascularized organ of the body and is a major site of vascular lesion development in HHT patients, more specifically in HHT2 patients (44). We next investigated whether vascular defects could be observed in the liver of the P9 tBMP9/10ib mice. Vascular pathology induction and drug treatments in pups were performed as above (Figure 2A). Latex dye tissue invasion was enhanced in the tBMP9/10ib liver and revealed a significant enlargement of the hepatic vessels (Figures 5A, 5B, and 5D). Hematoxylin and eosin (H&E) staining showed the presence of marked local liver injury, characterized by the presence of significant hepatocyte vacuolation and hepatocellular necrosis (Figures 5E, 5F, 5H, 5I, and S7), a pathology that could result from ischemic events.

**Figure 5.**
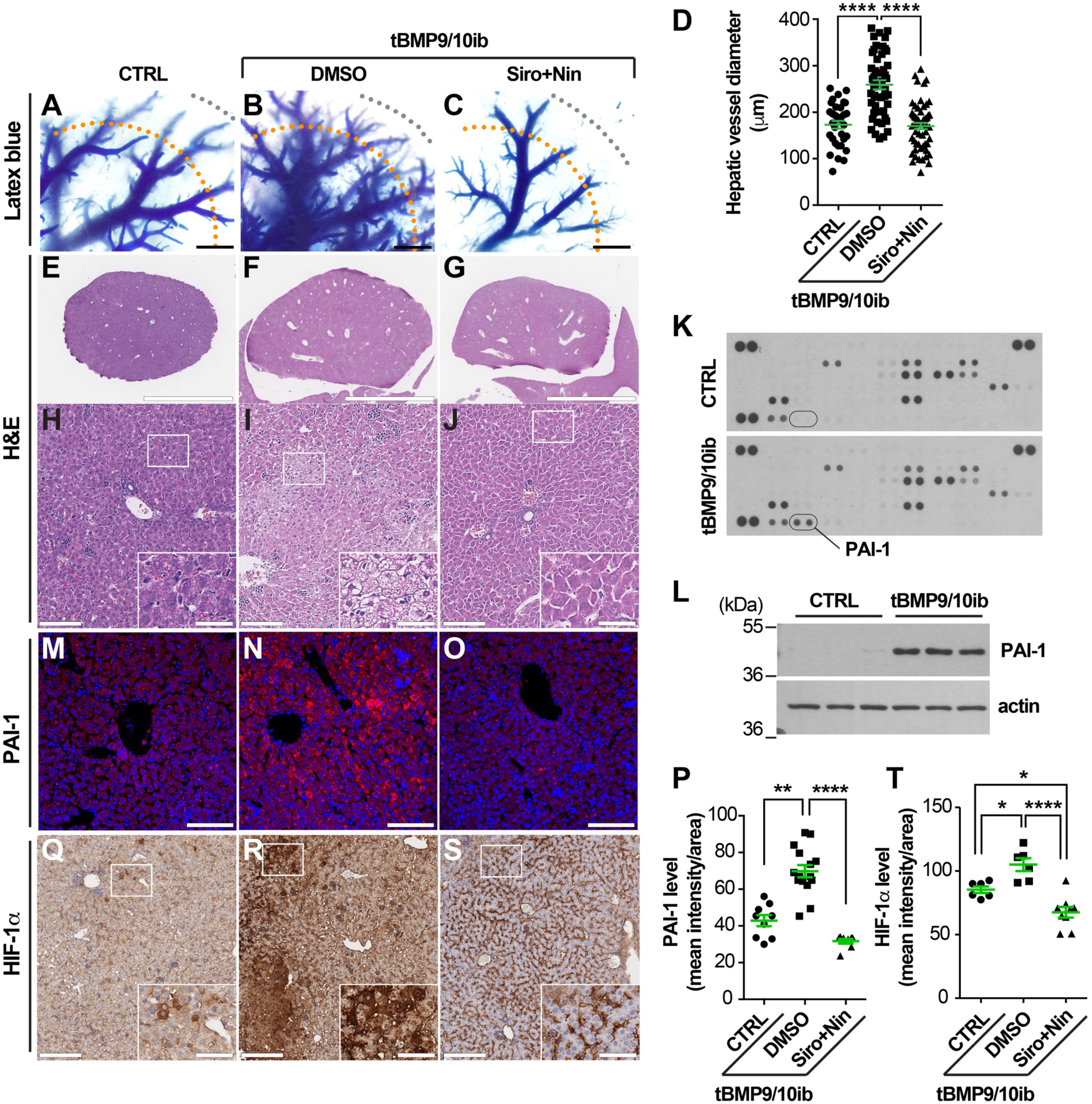
Liver vascular pathology and liver disease in tBMP9/10ib mice: Effects of Siro+Nin treatment. **(A-C)** Latex blue dye-perfused livers from tBMP9/10ib mice treated with vehicle (DMSO, B) or Siro+Nin (C), and control mice treated with DMSO (CTRL, A). **(D)** Scatter plot measuring hepatic vessel diameter. Diameters of these vessels (most likely arterial) were measured at 2,000 µm (orange dotted lines) from the margin of the liver (grey dotted lines). Data represent mean ± SEM (n = 4 livers per condition); Kruskal-Wallis test, post hoc Dunn’s multiple comparison test. **(E-J)** Liver sections stained with H&E at low magnification (E-G) and high magnification (H-J) from tBMP9/10ib and control mice treated as in (A-C), insets in (H-J) indicate magnified areas. **(K-O)** PAI-1 immunodetection by proteomic array (K, pooled homogenates from n = 3 mice) and WB (L, n = 3 mice) of liver homogenates, and by immunofluorescence staining in liver sections (M-O) from mice treated as in (A-C). **(P)** Scatter plot measuring PAI-1 levels, expressed as a mean intensity of fluorescence per sampled area. Data represent mean ± SEM (n = 3 livers per condition); Kruskal-Wallis test, post hoc Dunn’s multiple comparison test. **(Q-S)** HIF-1α IHC in liver sections from mice treated as in (A-C), insets indicate magnified areas. Histology data are representative of at least 3 independent mice/condition. Scale bars, 1000 µm (A-C), 3 mm (E-G), 100 µm (H-J, M-O, Q-S), 35 µm in insets (H-J, Q-S). **(T)** Scatter plot measuring HIF-1α levels, expressed as a mean intensity of fluorescence per sampled area. Data represent mean ± SEM (n = 3 livers per condition); one-way ANOVA test, post hoc Tukey’s multiple comparison test. \**P*<0.05, \*\**P*<0.01, \*\*\*\**P*<0.0001.

To gain insight into the mechanism of liver injury, we performed a proteome array for angiogenesis-related factors. This screen identified plasminogen activator inhibitor 1 (PAI-1) as a protein strongly upregulated in the tBMP9/10ib liver, compared to livers from control mice (Figure 5K). PAI-1 is of interest because it is upregulated during hypoxia (45), and might thus represent a response to ischemia. PAI-1 elevation in the tBMP9/10ib liver was confirmed by Western blot (WB, Figure 5L) and IHC (Figures 5M, 5N, and 5P) analyses. To verify that hypoxia is occurring in the tBMP9/10ib liver, tissue sections were stained for hypoxia-inducible factor-1α (HIF-1α), a transcriptional factor marker and master regulator of hypoxia (46). We found a strong upregulation of liver HIF-1α expression in tBMP9/10ib mice, compared to control mice (Figures 5Q, 5R, and 5T). These data show that tBMP9/10ib mice develop a robust vascular pathology in the liver. Strikingly, Siro+Nin treatment of the tBMP9/10ib mice significantly prevented the hyperproliferative vascular pathology and vessel dilation phenotype of the liver (Figures 5A-5D). In addition, Siro+Nin efficiently reduced hepatocyte vacuolation and necrosis (Figures 5E-5J and S7), as well as prevented the overexpression of PAI-1 and HIF-1α in the liver of the tBMP9/10ib mice (Figures 5M-5T). Together, these results demonstrate that Siro+Nin combination prevented vascular pathology in the liver, as well as blocked liver disease in tBMP9/10ib mice.

To further test the therapeutic potential of the Siro+Nin combination, we determined: (1) whether gene expression changes could be detected in the tBMP9/10ib whole-liver tissue and (2) whether treatments with the two drugs (administered alone or in combination) could correct these transcriptomic changes. When we plotted the transcript expression changes (measured as log fold change values, logFC) in tBMP9/10ib pups vs. normal controls, against the transcript expression logFCs obtained after treating the tBMP9/10ib pups with Siro+Nin vs. vehicle (DMSO), an inverse correlation was found for Siro+Nin treatment (r^2^ = 0.380, *P*<0.001; Figure 6A). An inverse correlation indicates that Siro+Nin treatment normalized some of the changes observed in the disease model (tBMP9/10ib pups) compared to normal controls. When the same comparison was done for Siro and Nin administered alone, a weaker correlation was measured (r^2^ = 0.193 and 0.190, respectively, *P*<0.001; Figure 6A). These results indicate that Siro+Nin combination better corrected the expression changes of liver transcripts detected in the tBMP9/10ib mice, than either drug administered alone.

**Figure 6.**
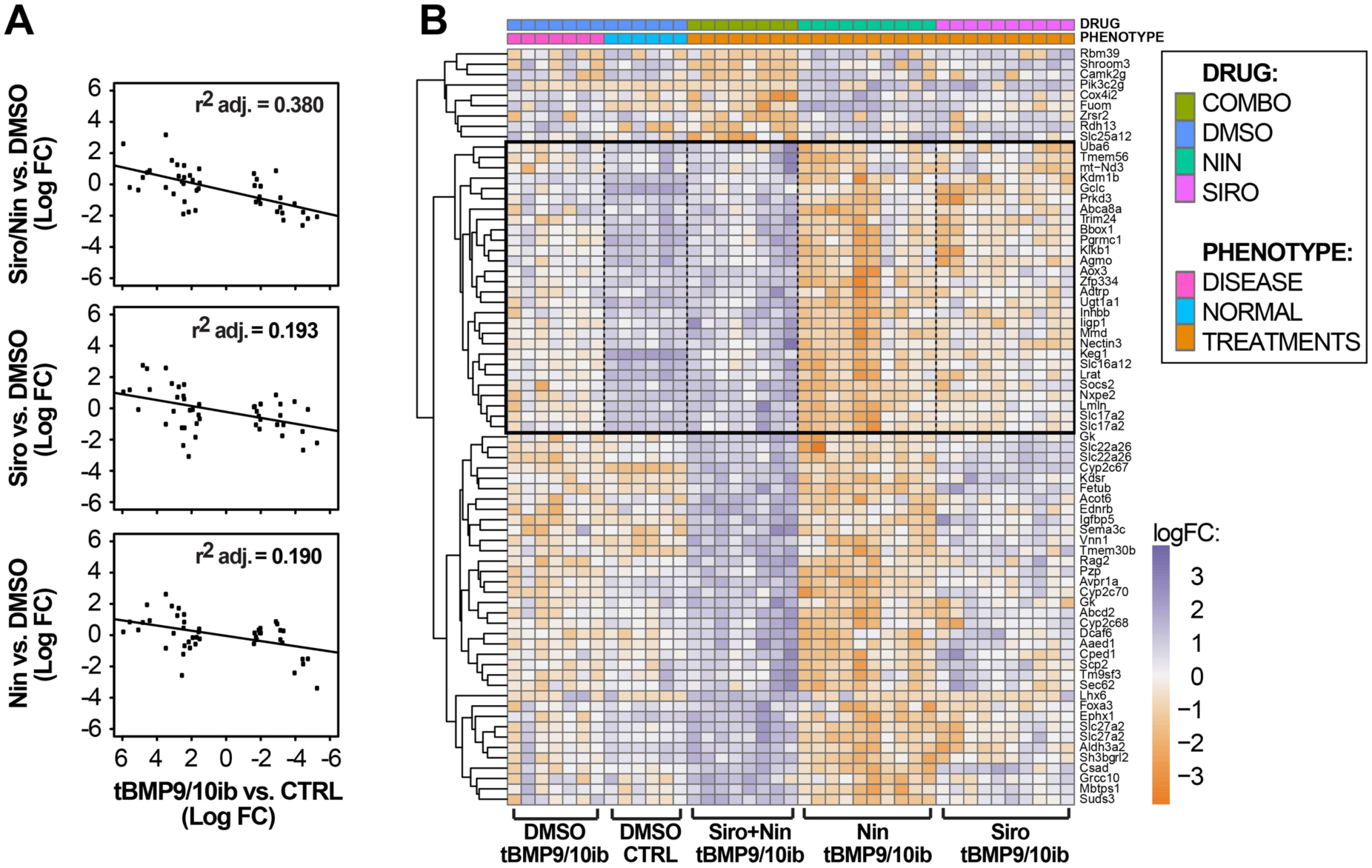
Siro+Nin corrects a gene expression signature detected in the tBMP9/10ib mouse liver. **(A)** Scatter plots comparing the liver gene expression changes (logFC, log fold change) observed upon treatment with Siro+Nin or individual drug (Siro or Nin), to the liver gene expression logFCs observed between tBMP9/10ib and normal control (CTRL) mice. The negative slope of the plots indicates that each treatment contributed to normalize the liver gene expression changes seen in tBMP9/10ib vs. CTRL mice. Correlation coefficients (r^2^) indicate a more robust normalization with the drug combination than with individual drugs. **(B)** Heatmap of liver transcripts synergistically differentially expressed by the drug combination. We identified the genes that responded to the drug combination differently than would be expected assuming an additive effect of both drugs. Non-interacting drug would produce a COMBO expression equal to the average of gene expression measured with each treatment (NIN or SIRO). The significant effects that we detected, measured with the statistical contrast COMBO-(NIN+SIRO)/2, produced gene expression changes that deviate from this average independent effect expectation. Transcripts were selected with FDR≤0.5%, log average expression >-5. For reference, liver transcripts in CTRL mice are shown on the heatmap. Gene expression in CTRL mice were used for normalization, but were not used in the selected transcripts that displayed synergistic drug response.

To illustrate the synergistic effect of the Siro and Nin treatments on transcript expression, we identified transcripts that were differentially changed by the drug treatments in the tBMP9/10ib liver (n = 6-9 biological replicates/group, false discovery rate (FDR) ≤0.5%, log average expression >-5; Figure 6B). These transcripts are shown in Figure S8 and Table S1. The black box in the heatmap of Figure 6B shows a subset of transcripts that were: (1) significantly changed by Siro+Nin, (2) not changed by either Nin or Siro administered in isolation, and (3) whose expression was normalized, as compared to normal controls. This subset of transcripts confirms that a synergistic effect exists between Siro and Nin treatments at the gene expression level, and that this effect could normalize a subset of deregulated transcripts in the tBMP9/10ib liver. A function enrichment analysis of the identified subset of normalized transcripts revealed a significant network of genes involved in cell and protein metabolism (e.g., *Uba6, Tmem56, mt-Nd3, Kdm1b, Gclc, Prkd3*; GeneMANIA (47)), a response indicative of strong gene expression changes consecutive to liver injury and changes in cell homeostasis and cell stress.

Together, these transcriptomic data show that Siro and Nin treatments synergized and partially opposed a deregulated gene expression signature detected in the tBMP9/10ib liver. These results are important because they confirm the interaction and combination efficacy of the two drugs in reducing overall vascular pathology in tBMP9/10ib mice.

### Siro+Nin reduces GI bleeding and anemia in adult Alk1 iKO mice

The effect of the drug combination was evaluated in an adult *Alk1* inducible knockout (iKO) mouse model (R26^CreER/+^;*Alk1*^2f/2f^). In this model, ALK1 deficiency is induced by tamoxifen administration to generate severe gastrointestinal (GI) bleeding and anemia in 9 days [Figure 7 and Refs. (39, 48)]. Daily treatment with the same doses of Siro+Nin used for the tBMP9/10ib mice (0.5 mg/kg Siro and 0.3 mg/kg Nin, i.p.), starting at the time of tamoxifen injection, significantly reduced GI bleeding (Figures 7A-7D) and significantly increased hematocrit level, RBC number, and hemoglobin level (Figures 7E-7G), compared to the vehicle-treated *Alk1* iKO controls.

**Figure 7.**
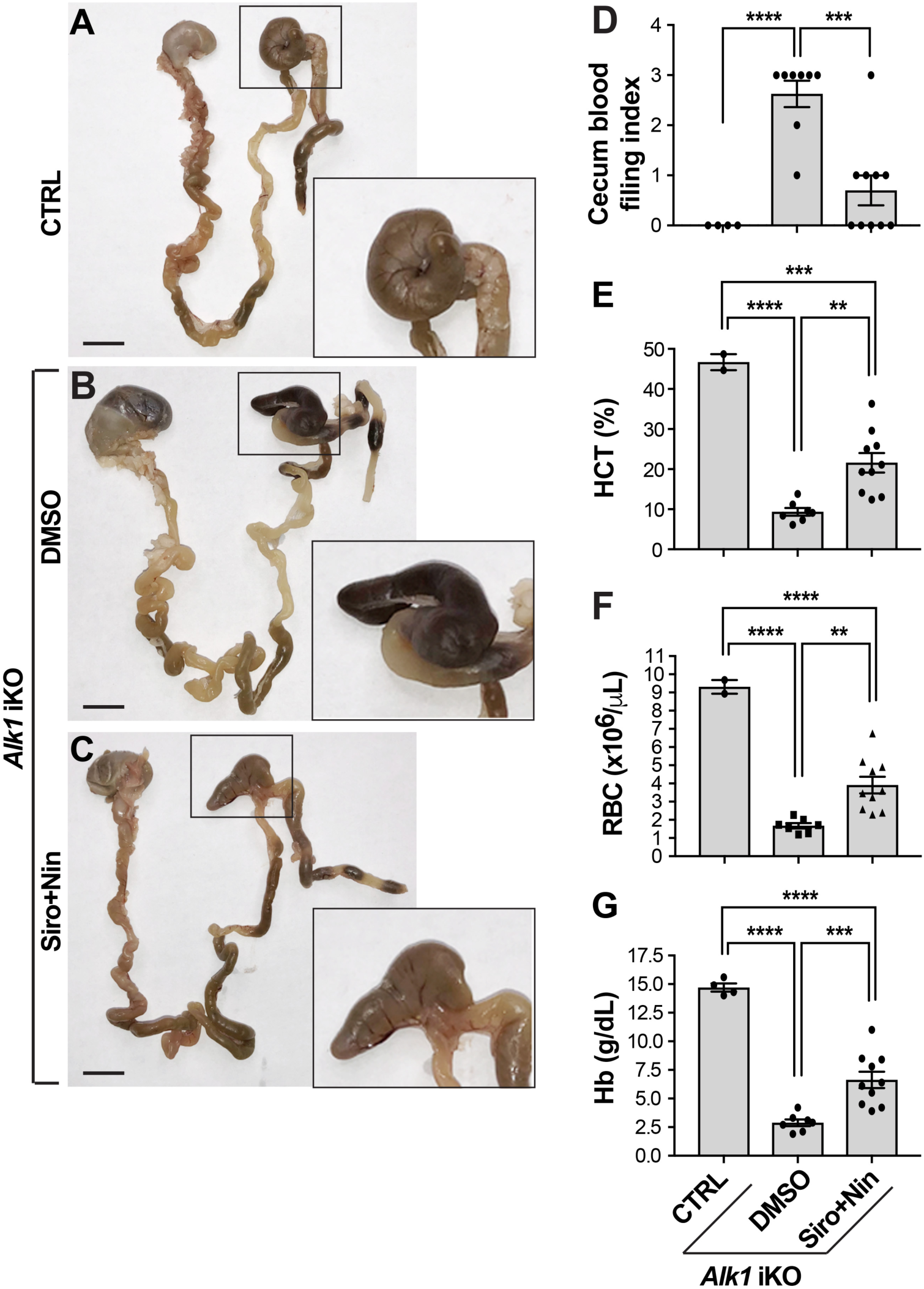
Siro+Nin effectively reduces GI bleeding and ameliorates anemia in adult *Alk1* iKO mice. **(A-C)** Gastrointestinal (GI) tract from stomach to rectum of 2-4 month-old CreER-negative control (CTRL, A) and *Alk1* iKO mice treated with vehicle (DMSO, B) or Siro+Nin (C). Higher magnifications show the cecum. Scale bars, 10 mm (A-C). **(D-G)** Scatter plots measuring cecum bleeding index (D), hematocrit (HCT) level (E), red blood cell (RBC) number (F), and hemoglobin (Hb) level (G). Data represent mean ± SEM (n = 2-4, 7-8, 10 mice for the CTRL, DMSO, and Siro+Nin groups, respectively); one-way ANOVA (E and F) and two-way ANOVA (G), Tukey’s multiple comparison test; \*\**P*<0.01, \*\*\**P*<0.001, \*\*\*\**P*<0.0001.

### In tBMP9/10ib mice, Siro and Nin prevent endothelial overactivation of mTOR and VEGFR2, respectively

We next assessed mTOR signaling in tBMP9/10ib mice by measuring the levels of phospho-mTOR (p-mTOR) and phospho-S6 (p-S6) in the liver and retina. WB analyses revealed robust increases in p-mTOR and p-S6 levels in whole-liver homogenates isolated from tBMP9/10ib mice, compared to control mice (Figure 8A). Treatment with Siro (alone or in combination with Nin) blocked S6 phosphorylation and normalized p-mTOR levels in the tBMP9/10ib mouse liver (Figures 8A and S9). Further examination of the tBMP9/10ib retinal tissue confirmed the presence of strong p-S6 immunoreactivity in the AVMs, which could significantly be blocked by Siro treatment of the mice (Figures 8B and 8C).

**Figure 8.**
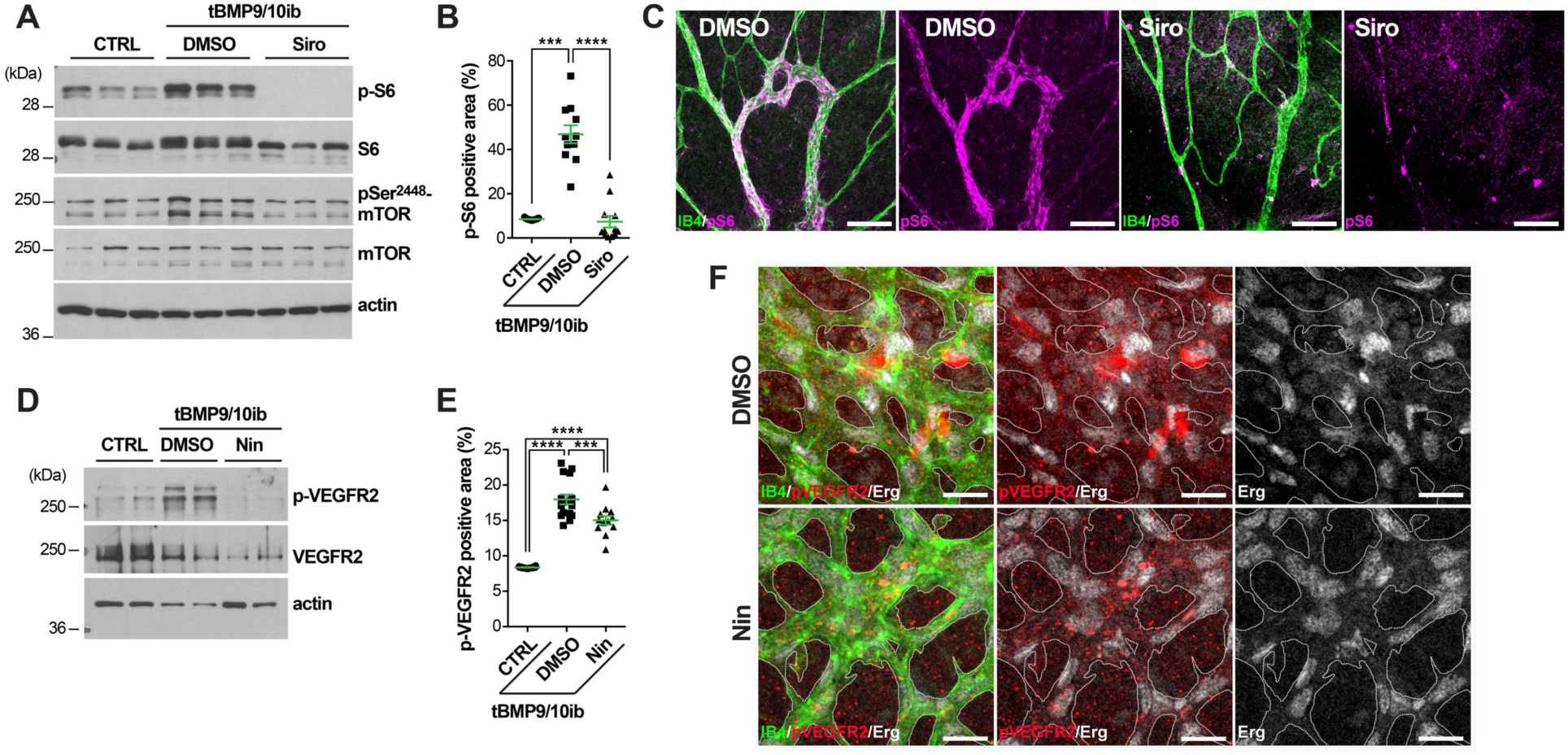
In tBMP9/10ib mice, Siro and Nin prevent endothelial overactivation of mTOR and VEGFR2, respectively. **(A-F)** Control (CTRL) and tBMP9/10ib mice were treated with DMSO and Siro (A-C) or Nin (D-F). Protein homogenates of whole liver from P6 mice (A) and of liver ECs isolated from P9 mice (D) were analyzed by WB using antibodies directed against the indicated proteins. Scatter plots show p-S6 (B) and p-VEGFR2 (E) positive area analyzed by IHC in the retina of P6 CTRL and tBMP9/10ib mice treated with DMSO and Siro (B) or Nin (E), expressed as a percentage of the retinal vascular area occupied by immunofluorescence staining. Data represent mean ± SEM (n = 4, 3, 3 mice for the CTRL, DMSO, and Siro (B) or Nin (E) groups, respectively); p-S6 analysis: Kruskal-Wallis test, post hoc Dunn’s multiple comparison test; p-VEGFR2 analysis: one-way ANOVA, Tukey’s multiple comparison test; \*\*\**P*<0.001, \*\*\*\**P*<0.0001. Representative immunofluorescence images show P6 retinas from tBMP9/10ib mice treated as in (B and E) stained with fluorescent isolectin B4 [green, (C and F)], anti-p-S6 [magenta, (C)], anti-p-VEGFR2 [red, (F)], and anti-Erg [white, (F)] antibodies. Scale bars, 100 µm (C) and 50 µm (F).

A significant elevation of activated phospho-VEGFR2 (p-VEGFR2) was detected in protein homogenates of liver ECs isolated from tBMP9/10ib mice (Figures 8D and S10) and in the tBMP9/10ib mouse retina (Figure 8E), which was significantly inhibited by Nin treatment of the mice (Figures 8D-8F). We also verified that drug combination did not interfere with the inhibitory effect of Nin on p-VEGFR2. As observed upon treatment with Nin alone, Siro+Nin fully inhibited VEGF-induced VEGFR2 activation in primary ECs (Figure S2). Altogether, these data confirmed that endothelial mTOR and VEGFR2 are overactivated in tBMP9/10ib mice and that Siro and Nin can respectively and efficiently block these signaling deregulations in vivo.

### Siro rescues Smad1/5/8 signaling by activating ALK2

Because the possibility that Siro might both prevent mTOR overactivation and rescue Smad1/5/8 signaling in HHT mice is appealing, we revisited the effect of Siro on Smad1/5/8 signaling in cell culture systems. We found that Siro increased Smad1/5/8 phosphorylation (p-Smad1/5/8) in a dose-dependent manner and increased the levels of the Smad1/5/8 downstream target ID1 (inhibitor of differentiation 1) in C2C12 myoblast cells (Figure S11A) and in human umbilical vein ECs (HUVECs, Figure 9A), at doses very comparable to Tac (32). A pharmacological approach using the pan-ALK inhibitor LDN-193189 (Figure 9B) and gene silencing (Figures 9C and S11B) in HUVECs further showed that ALK2 is required for the stimulatory effect of Siro on Smad1/5/8 signaling.

**Figure 9.**
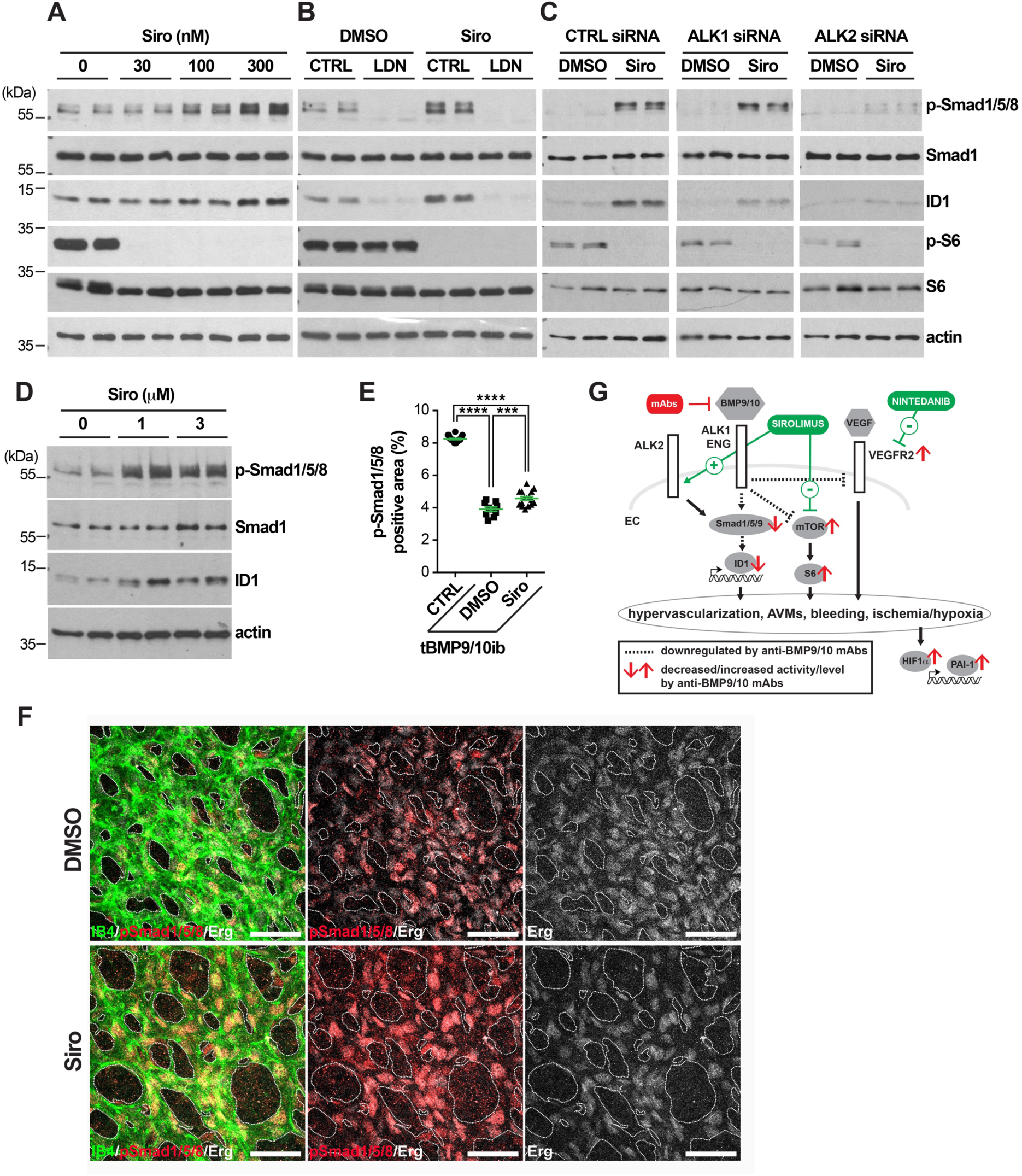
Siro rescues Smad1/5/8 signaling by activating ALK2. **(A-D)** HUVECs (A-C) and HHT2 patient-derived BOECs (D) were treated for 3h with vehicle (DMSO) or Siro in complete medium (conditioned for 2-3 days) and at the indicated concentrations (A and D) or at 300 nM (B and C), in the absence or presence of LDN-193189 [LDN, 1 µM, (B)], or the indicated siRNA treatments (C). Cell extracts were analyzed by WB using antibodies directed against the indicated proteins. **(E)** Scatter plot measuring p-Smad1/5/8 positive area in the retina of P6 control (CTRL) and tBMP9/10ib mice treated with DMSO or Siro, expressed as a percentage of the retinal vascular area occupied by immunostaining. Data represent mean ± SEM (n = 4, 3, 3 mice for the CTRL, DMSO, and Siro groups, respectively); \*\*\**P*<0.001, \*\*\*\**P*<0.0001; one-way ANOVA, Tukey’s multiple comparison test. **(F)** Representative images showing P6 retinas from tBMP9/10ib mice treated as in (E) stained with fluorescent isolectin B4 (green), anti-Erg (white), and anti-p-Smad1/5/8 (red). Scale bars, 50 µm. **(G)** Schematic illustration of the proposed synergistic effects of Siro and Nin on endothelial Smad1/5/8, mTOR, and VEGFR2 signaling pathways.

To further demonstrate efficacy in a cell culture system more directly relevant to the pathophysiology of human HHT, we used blood outgrowth ECs (BOECs) isolated from a HHT2 patient genetically confirmed to carry the disease-causing ALK1 T372fsX truncation (32). Treating these HHT2 patient BOECs with Siro also increased p-Smad1/5/8 and ID1 levels (Figure 9D). Finally, IHC analyses showed that in vivo Siro was able to modestly but significantly rescue the decrease of p-Smad1/5/8 in the retinal vasculature (Figures 9E and 9F). Thus, Siro displayed potent in vitro Smad1/5/8 signaling activating properties in EC cultures, including in HHT patient-derived ECs, and significantly rescued Smad1/5/8 loss-of-function in vivo in tBMP9/10ib mice.

## DISCUSSION

Major progress in techniques of interventional radiology, embolization, and cauterization have considerably reduced the risk of hemorrhage in HHT patients. However, the recurring and progressive nature of the disease continues to make HHT a highly debilitating disorder associated with life-threatening sequelae. Several disease-modifying therapies have been proposed in pre-clinical models and some of them are currently being investigated in clinical trials (49). Notably, Tac was found to be associated with Smad1/5/8 activating properties in ECs and showed some efficacy in reducing pathology in HHT (32) and PAH (37) models and patients (50–52). Here we propose that Siro, a structural analog of Tac and inhibitor of mTOR, is a more potent blocker of AVM development in HHT mice. We found that Siro acts as a dual modulator by stimulating Smad1/5/8 while inhibiting mTOR, in both cell cultures and HHT mice, thus correcting two key signaling defects of the HHT ECs.

In cell cultures, Tac was proposed to activate Smad1/5 by facilitating dissociation of the binding repressor FKBP12 from ALK1, ALK2, and ALK3, while Siro only recruited ALK2 (37). Here we confirmed in HUVECs that Siro could activate endothelial Smad1/5/8 by specifically stimulating ALK2. Although these data would need to be confirmed in vivo, they suggest that the intriguing possibility that ALK2 activation might compensate for ALK1 loss-of-function. Although attractive from a mechanistic aspect, this mechanism might be risky as ALK2 activating mutations are associated with fibrodysplasia ossificans progressiva, a disorder characterized by progressive heterotopic bone formation in muscle tissues (53). However, Siro, as well as Tac that also activates ALK2, are not known to lead to any adverse effects associated with mechanisms of abnormal ossification in humans. In addition, Siro showed a potent anti-AVM effect at doses (9-22 nM in mouse serum) comparable to its use in the clinical setting (whole-blood trough concentrations of ∼15-20 nM for prophylaxis of transplant rejection, for instance).

Our data highlight a beneficial interaction and synergy between Siro and Nin treatments in HHT mice. This interaction was observed by RNA-Seq of the tBMP9/10ib liver, which revealed the presence of deregulated genes that were fully normalized only when Siro and Nin were co-administered. This transcriptional interaction translated into a robust synergy of the two drug treatments and a significant improvement of the Siro effect at preventing AVMs in HHT mice. We found that Nin acted, at least in part, by blocking the overactivation of VEGFR2 in ECs of the tBMP9/10ib mice. Nin is not specific to VEGFRs, as it also targets FGFRs and PDGFRs, two other receptor families involved in angiogenesis (54). Further investigation will be needed to determine whether interference of these other RTK receptor pathways might be involved in the anti-AVM effect of the Siro+Nin combination. Previous work suggests, however, that this might not be the case for PDGFRs, since PDGF signaling activation (not inhibition) was found to mitigate vessel wall defects in an *Eng* KO mouse model (55).

The current data provide further evidence that HHT pathogenesis requires the overactivation of the mTOR and VEGFR2 pathways (Figure 9G). How Smad1/5/8, mTOR, and VEGFR2 control each other during the disease process remains unclear. Although loss-of-function mutations in BMP9-ALK1-endoglin-Smad4 signaling are necessary to cause HHT, they are not sufficient. Indeed, additional genetic, environmental, and/or developmental modifiers are required for AVM development, a process referred to as the double/multiple hit hypothesis (56). Notably, VEGF stimulation or wounding, two triggers of angiogenesis, were reported to be required to generate brain and skin AVMs in adult ALK1 deficient mice (43). Furthermore, in HHT neonate mouse models, including in tBMP9/10ib pups, it is believed that retinal AVMs develop spontaneously and aggressively because vessel development is actively occurring through angiogenesis. Synergy between BMP9/10 blocking and VEGFR2 activation during the normal process of angiogenic vascular development would thus provide in these models the ideal milieu for lesion development. Therefore, although there is now strong evidence that Smad1/5/8 reduction overactivates VEGF signaling to cause vascular pathology, these two pathways are initiated by independent triggers. Our data showing that Nin and Siro treatments have clear synergistic effects support this notion and further show that the two pathways can independently signal in ECs to cause AVMs.

Recent studies have demonstrated that PI3K-Akt signaling is overactivated in several HHT models and that its inhibition reduces the AVMs (25, 30, 31). In line with this observation, we show here that, in the tBMP9/10ib liver, there is an increase of p-Ser^2448^-mTOR, a phosphorylation mostly associated with mTORC1, the mTOR complex downstream from PI3K-Akt, which is directly targeted by Siro (57). In addition, Siro treatment, which fully prevented p-Ser^2448^-mTOR and S6 overactivation, reduced AVM number in our model, suggesting that mTOR, most likely downstream from PI3K-Akt, is a major player in AVM development. We further found that, although Siro fully inhibited mTOR overactivation in vivo, its anti-AVM effect was only partial and could be significantly potentiated upon VEGFR2 inhibition by Nin, indicating that VEGFR2 can signal independently of the PI3K-Akt-mTOR pathway to control the AVMs. This observation concurs with recent data obtained in Smad4 deficient mice that indicate that abnormal PI3K-Akt activation in ECs is primary controlled by blood flow and not VEGFR2 (24, 58).

An important question relates to the drug dosing used in this study and whether it allows the administered drugs to reach concentrations within the expected therapeutic range. We measured in average 9.0 nM Siro and 5.3 nM Nin at P6, and an increase to 22.6 nM Siro and 24.7 nM Nin at P9 in the pup serum. Previously published in vitro experiments reported a Siro IC50 of ∼0.1 nM for mTOR (59) and a Nin IC50 of ∼13 nM for VEGFR2 (60). At the tested dosing, circulating concentrations of Siro and Nin are therefore expected to inhibit in vivo mTOR and VEGFR2, which is in line with the observed effects on these targets in tBMP9/10ib mice (Figure 8). For Smad1/5/8 stimulation by Siro, our experiments in HUVECs showed that the lowest effective concentration is 100 nM (Figure 9A), which is about 4 times higher than the concentration detected in the serum. We found nonetheless that Siro treatment modestly but significantly increased p-Smad1/5/8 in the mouse retina, indicating that the drug was delivered in sufficient quantities to stimulate Smad1/5/8 signaling in vivo (Figure 9). We have not obtained clear experimental data that explain this apparent discrepancy. However, it is important to note that concentrations used in vitro and drug serum concentrations are difficult to compare. Indeed, drug concentration measurements in the circulation do not consider the pharmacokinetic and bioavailability of the drugs. One important parameter is cellular uptake and accumulation, which can significantly reduce serum concentration while increasing local drug delivery. Altogether these data suggest that the administered drugs reach concentrations within the expected therapeutic range.

We found that Siro+Nin treatment prevented anemia in P9 tBMP9/10ib pups and adult *Alk1* iKO mice (Figures 2B and 7D-G). In tBMP9/10ib pups, drug treatment generated a supranormal elevation of the CBC parameters. In some models of anemia, Siro was reported to ameliorate anemia by increasing stress erythropoiesis (61). In this context, the supranormal elevation of the CBC parameters measured after drug treatment could be due, in part, to changes in erythropoiesis. To test this hypothesis, additional experiments were performed by using the recently described method of monitoring erythropoiesis using CD44/Ter119/FSC as markers of terminal erythroid differentiation (62). We measured a significant increase in the number of reticulocytes in the bone marrow, but not in the spleen, of Siro+Nin-treated tBMP9/10ib mice (Figure S12). These data suggest that although no stress erythropoiesis induction was observed, an increase in RBC production occurred and thus could explain, at least partly, the supranormal levels of hematocrit, RBCs, and hemoglobin measured after the drug treatment. Further investigation will be required to clearly define the origin of the observed effects, for instance, by looking at the earlier RBC precursors (62). Other experiments will also be needed to precisely determine how much of the corrected anemia is driven by a bleeding reduction (Figure 2K) vs. an increased RBC production (Figure S12) and whether the drug effect on RBC production could be related to developmental aspects in newborn mice. In support to the latter possibility, a supranormal elevation of the RBC counts was not found in treated adult *Alk1* iKO mice (Figure 7), further supporting our current interpretation that anemia correction is primarily driven by a bleeding reduction.

Prophylaxis of rejection in liver transplantation includes treatment with immunosuppressive drugs, primarily Tac and more rarely Siro. Recently, liver AVM pathology recurrence was reported in HHT patients who underwent liver transplantation, while they were treated with Tac (63). Our study suggests the possibility that prioritizing Siro vs. Tac (together or not with Nin) in the antirejection regimen might also help in specifically preventing spreading of vascular disease in healthy liver grafts.

More broadly, the beneficial effect of Siro and mTOR inhibition has already been described in models of pathological angiogenesis (64). In addition, Siro treatment has produced promising results in patients affected by vascular anomalies (65). Our observation that Siro and Nin mechanistically synergize to block the pathogenic process of HHT might thus have important ramifications for the treatment of other vascular disorders, such as the vascular anomaly diseases.

In conclusion, the present data show that Siro and Nin combination concurrently normalizes endothelial Smad1/5/8, mTOR, and VEGFR2 pathways to synergistically and efficiently oppose HHT pathogenesis and the associated vascular pathology in neonate and adult mouse models of the disease. We thus propose that Siro+Nin combination has therapeutic potential in HHT.

## METHODS

### Mice

This study was performed using C57BL/6 mice (The Jackson Laboratory). R26Cre^ER/+^;*Alk1*^2f/2f^ mice generation and *Alk1* deletion using tamoxifen were described previously (39, 48). Mice were maintained in regular housing conditions and were allowed free access to water and maintenance diet.

### Transmammary-delivered immunoblocking of BMP9 and BMP10, and drug treatments

tBMP9/10ib pups and age-matched control pups (treated with isotype controls) were generated as previously described (22, 32). Pups (litter size, n = 7-9 mice; average mouse weight, 2 grams) were treated with vehicle (0.1% DMSO in saline), Siro (0.5 mg/kg bw), and Nin (0.3 mg/kg bw) following the different combinations and schedules described in the Figures. Isotype control pups were injected with vehicle.

### RNA-Seq, read alignments, and gene expression differential expression analysis

RNA was processed for RNA-Seq at the Genomics Resources Core Facility, Weill Cornell Medical College, New York, NY, on an Illumina HiSeq 4000 instrument. Alignments of reads to the transcriptome were performed with Kallisto version 0.44.0. We aligned reads to the mouse transcriptome downloaded from Ensembl (Mus_musculus.GRCm38.cdna.all.fa.gz). Alignment was run on a MacOS laptop with 16GB of memory, using the NextflowWorkbench (66) to orchestrate alignment and count aggregation for the analyzed samples. RNA-Seq transcript counts were analyzed with analysis scripts written with the MetaR languages (67). Additional details of the analyses are described in *Supplemental Material*.

### Statistics

For multiple-group comparisons, a one-way ANOVA test was performed (when data fulfilled the assumption of normality for parametric tests, using a D’Agostino & Pearson omnibus test) and a post hoc Tukey’s multiple comparison test was used. When normality was not achieved, a non-parametric Kruskal-Wallis test and post hoc Dunn’s multiple comparison test were used. For two-group comparisons, when normality was not achieved, a Mann-Whitney test was performed instead. P values < 0.05 were considered statistically significant. Analyses were done using GraphPad Prism.

### Study approvals

Study subject (HHT2 patient carrying the ALK1 T372fsX mutation) provided voluntary and written informed consent using a form approved by The Feinstein Institutes for Medical Research Institutional Review Board (IRB). Study subject BOECs were isolated and cultured using a protocol approved by the Institute’s IRB. All animal procedures were performed in accordance with protocols approved by The Feinstein Institutes for Medical Research and Barrow Neurological Institute Institutional Animal Care and Use Committees, and conformed to the NIH Guide for the Care and Use of Laboratory Animals and ARRIVE guidelines.

## Additional Methods are described in Supplemental Material

### Author contributions

PM conceived the study. SR and PM designed and supervised the study. JP and LB performed the hematological studies. HC and SPO performed the experiments in *Alk1* iKO mice. LD and PW performed the heart function measurements. PKC and CNM provided the HUVECs. MH and YAA performed the LC-MS. FC performed the bioinformatics analyses. SR, HZ, PC, ANK, RP, and MG performed all the other experiments. SR, LB, FC, and PM wrote the manuscript. CNM and SPO edited the manuscript.

## Acknowledgments

This study was supported by NIH/NHLBI grants R01HL139778 (to PM) and R01HL128525 (to SPO), DOD/CDMRP grant PR161205 (to SPO and PM), a Feinstein Institutes for Medical Research fund (to PM), Leducq foundation (ATTRACT, to SPO), and Barrow Neurological Foundation (to SPO).

## SUPPLEMENTAL FIGURES

**Figure S1.**
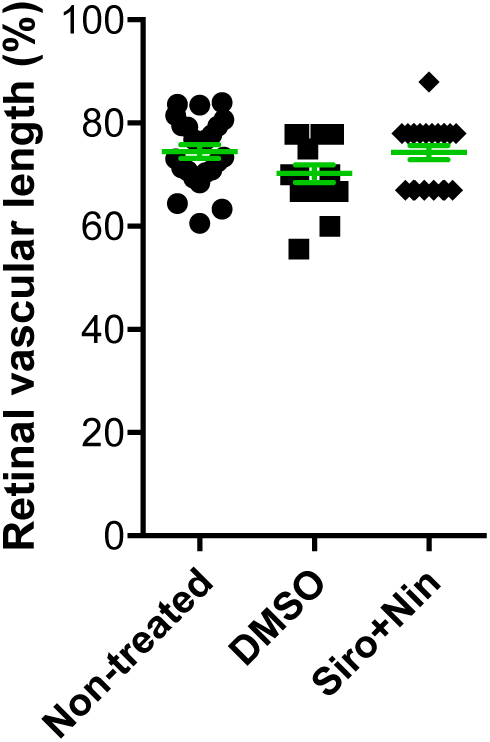
Siro+Nin does not affect normal vascular development in the mouse retina. Scatter plot shows retinal vascular length in P6 pups non-treated or treated with DMSO or Siro+Nin, as in Figure 1A. Data show the % of total retinal length occupied by the vasculature and represent mean ± SEM (n = 6, 4, 5 mice for the non-treated, DMSO, and Siro+Nin groups, respectively). One-way ANOVA and Tukey’s multiple comparison tests revealed no significant difference between groups.

**Figure S2.**
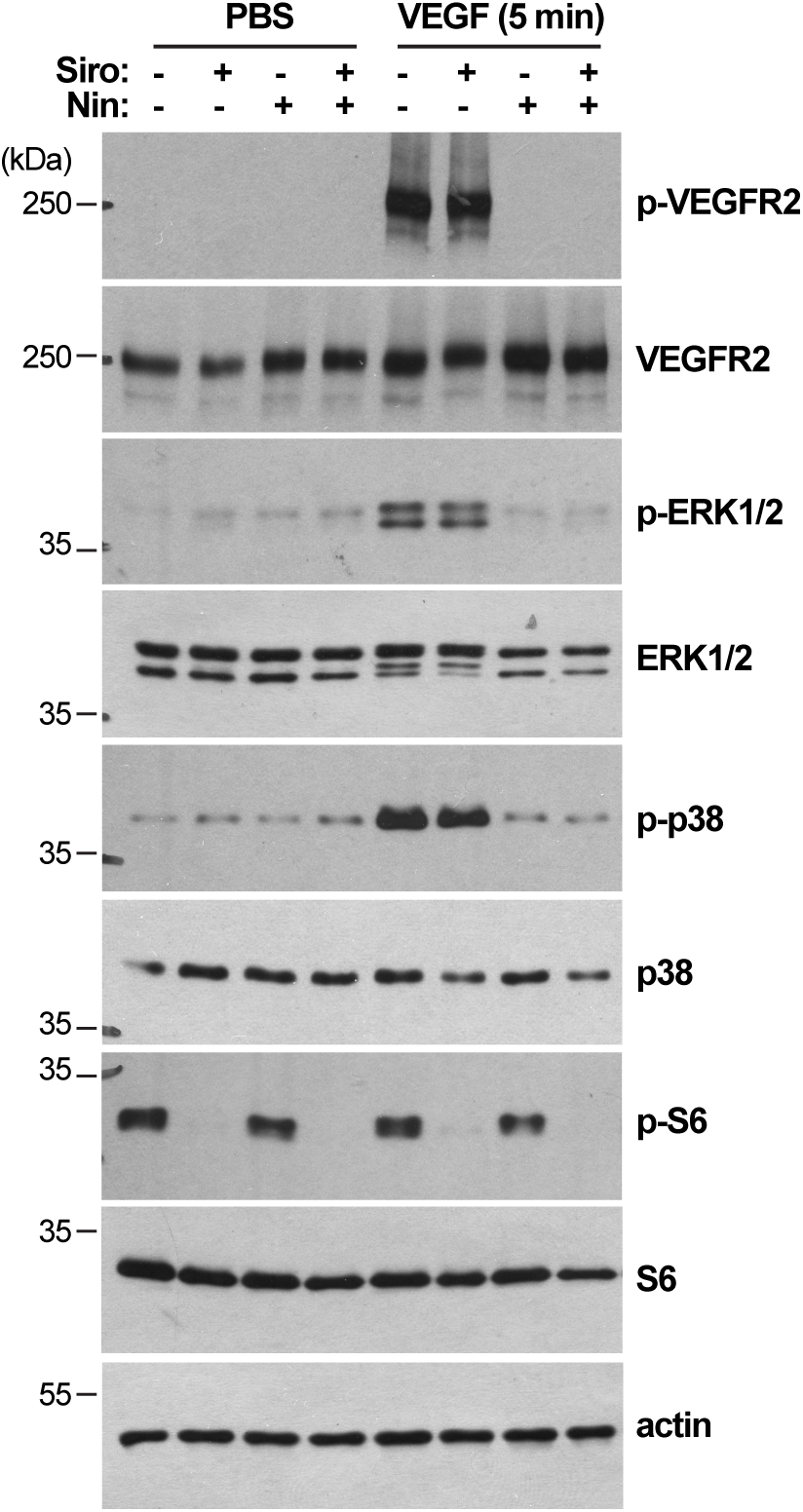
Siro does not inhibit VEGF-mediated ERK1/2 or p38 activation. HUVECs treated with vehicle [DMSO, (-)], Siro (300 nM), or Nin (300 nM), were stimulated for 5 min with VEGF (25 ng/mL), following a 24h starvation in 0.05% FBS-containing medium. Cell extracts were analyzed by WB using antibodies directed against the indicated proteins.

**Figure S3.**
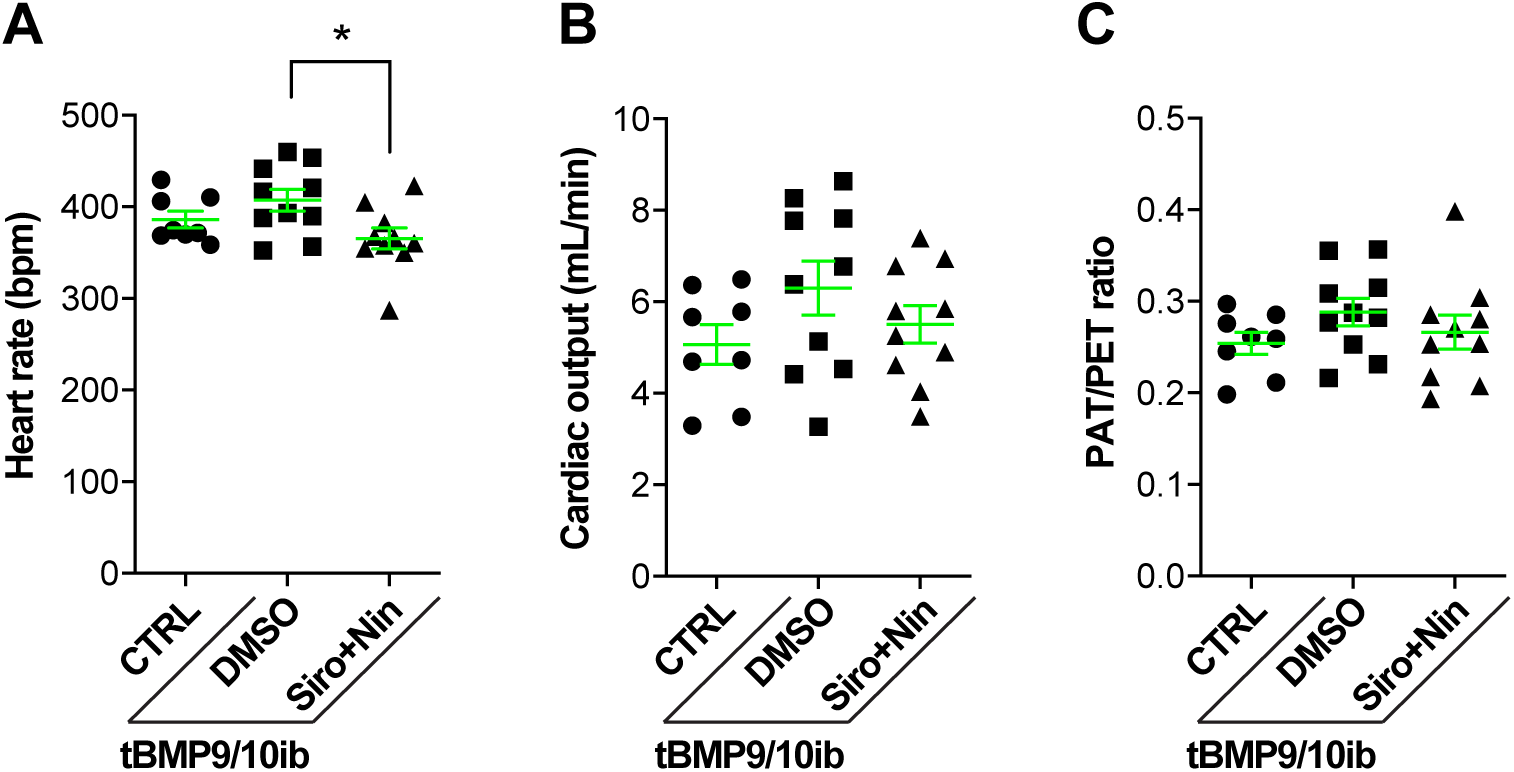
Siro+Nin does not affect cardiac function in tBMP9/10ib mice. Scatter plots show heart rate (A), cardiac output (B), and PAT/PET ratio (C) of P9 CTRL and tBMP9/10ib mice treated as in Figure 2A. Data represent mean ± SEM (n = 8, 10, 10 mice for the CTRL, DMSO, and Siro+Nin groups, respectively); one-way ANOVA, Tukey’s multiple comparison test; \**P*<0.05.

**Figure S4.**
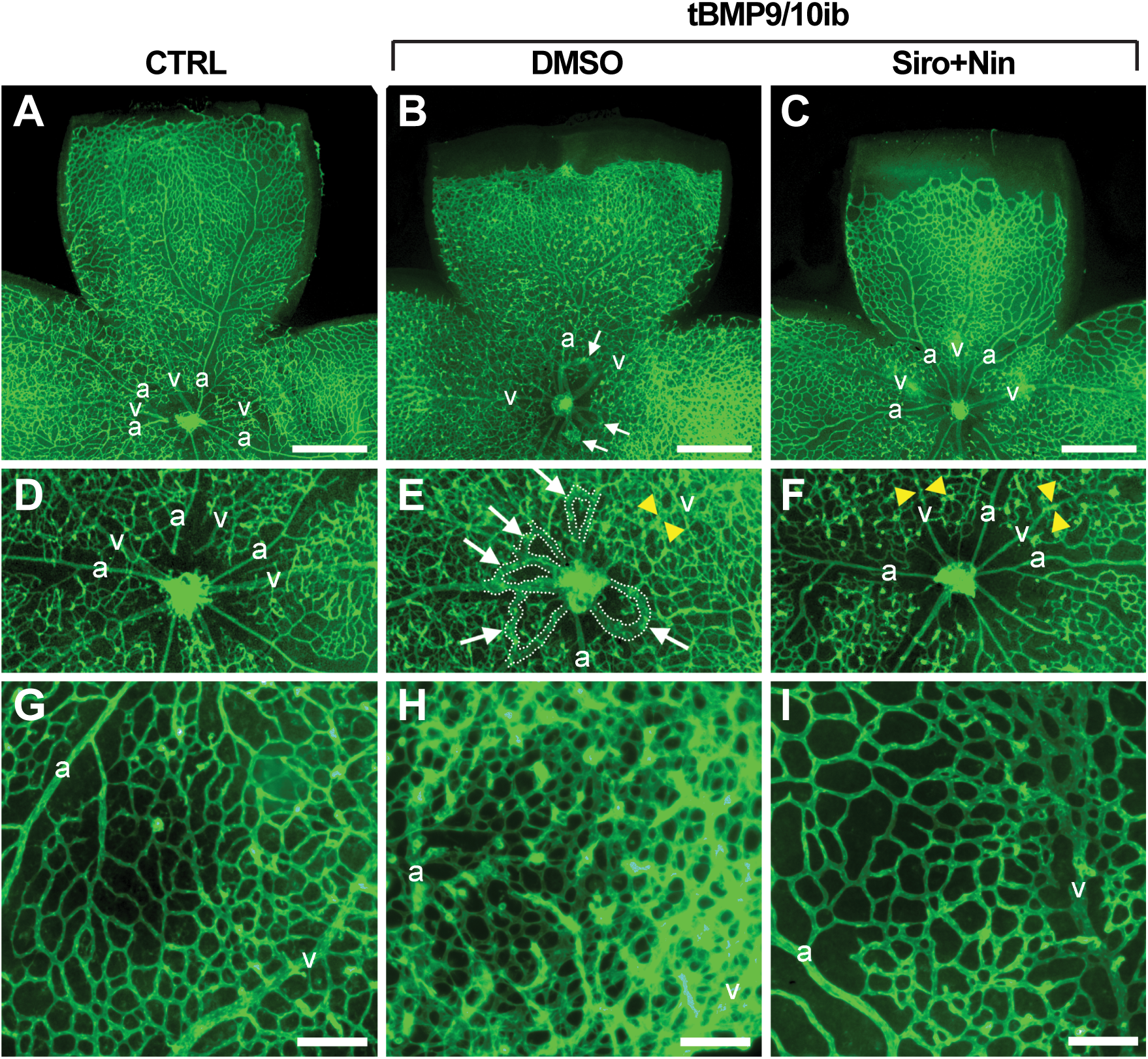
Siro+Nin prevents vein dilation, hypervascularization, and AVMs in the tBMP9/10ib mouse retina. **(A-I)** Representative images of retinas stained with fluorescent isolectin B4 (green) from P9 CTRL and tBMP9/10ib mice treated with DMSO and Siro+Nin, as in Figure 2A. Higher magnification in (D-F) shows retinal vein diameter [yellow arrowheads in (E and F) and retinal vascular fields (plexus area) between an artery (a) and a vein (v) (G-I). White arrows denote AVMs. Scale bars, 500 µm (A-C) and 100 µm (G-I).

**Figure S5.**
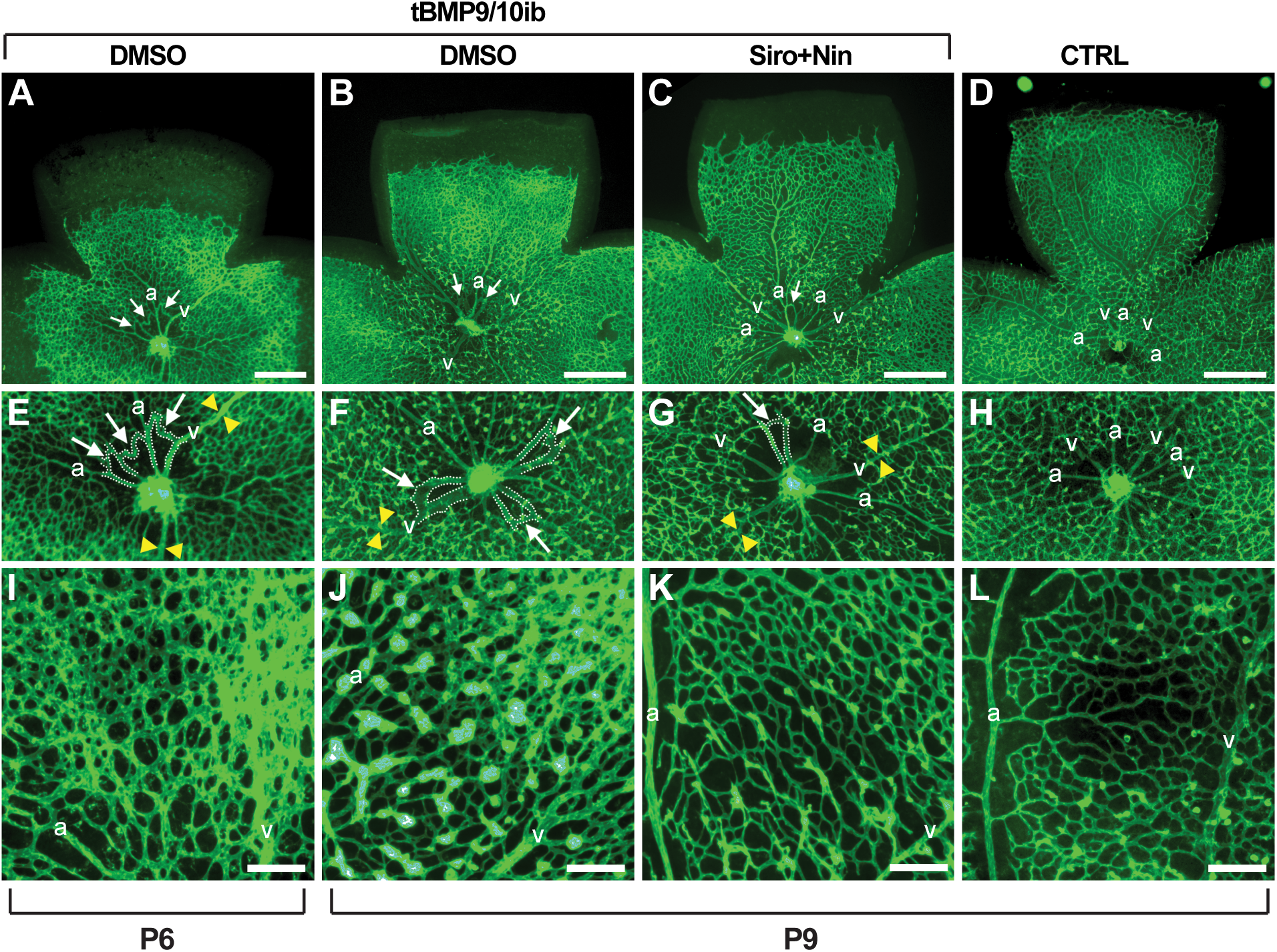
Siro+Nin reverses vein dilation, hypervascularization, and AVMs in the tBMP9/10ib mouse retina. **(A-L)** Representative images of retinas stained with fluorescent isolectin B4 (green) from CTRL and tBMP9/10ib mice (at the indicated postnatal day) treated with DMSO and Siro+Nin, as in Figure 3A. Higher magnification in (E-H) shows retinal vein diameter [yellow arrowheads in (E, F, and G)] and retinal vascular fields (plexus area) between an artery (a) and a vein (v) (I-L). White arrows denote AVMs. Scale bars, 500 µm (A-D) and 100 µm (I-L).

**Figure S6.**
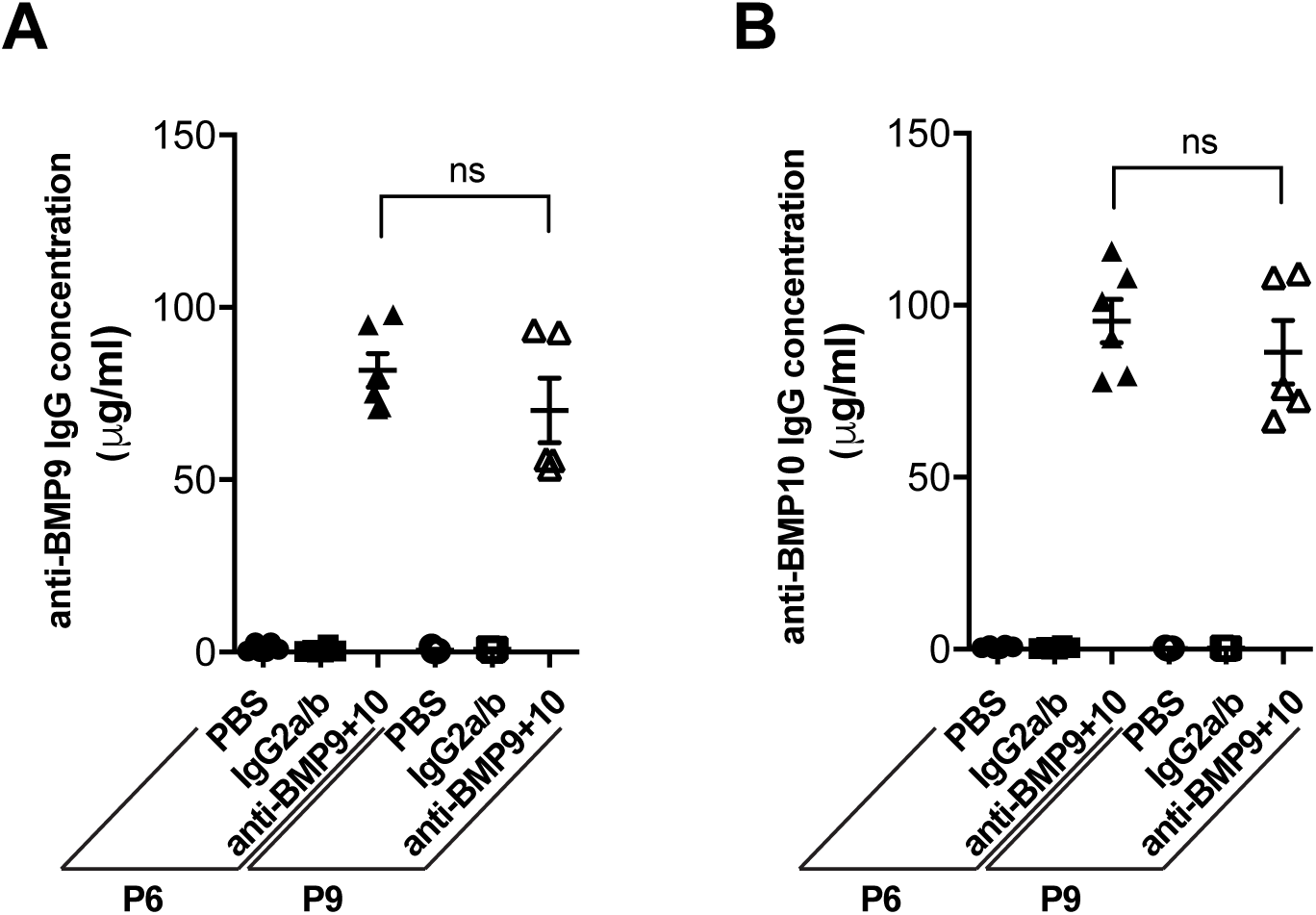
Transmammary-delivered BMP9 and BMP10 blocking antibodies persist in the circulation of mouse neonates. ELISAs were performed to measure anti-BMP9 **(A)** and anti-BMP10 **(B)** IgG concentration in the serum of P6 and P9 pups treated either at P3-P5 or P3-P8 through lactation from dams injected i.p. with PBS, isotype controls (IgG2a/b), or anti-BMP9 + anti-BMP10 antibodies (anti-BMP9+10). Data represent mean ± SEM (n = 6 and 5 mice for the P6 and P9 groups, respectively). One-way ANOVA and Tukey’s multiple comparison tests revealed no significant difference between P6 and P9 mice for the anti-BMP9+10 groups.

**Figure S7.**
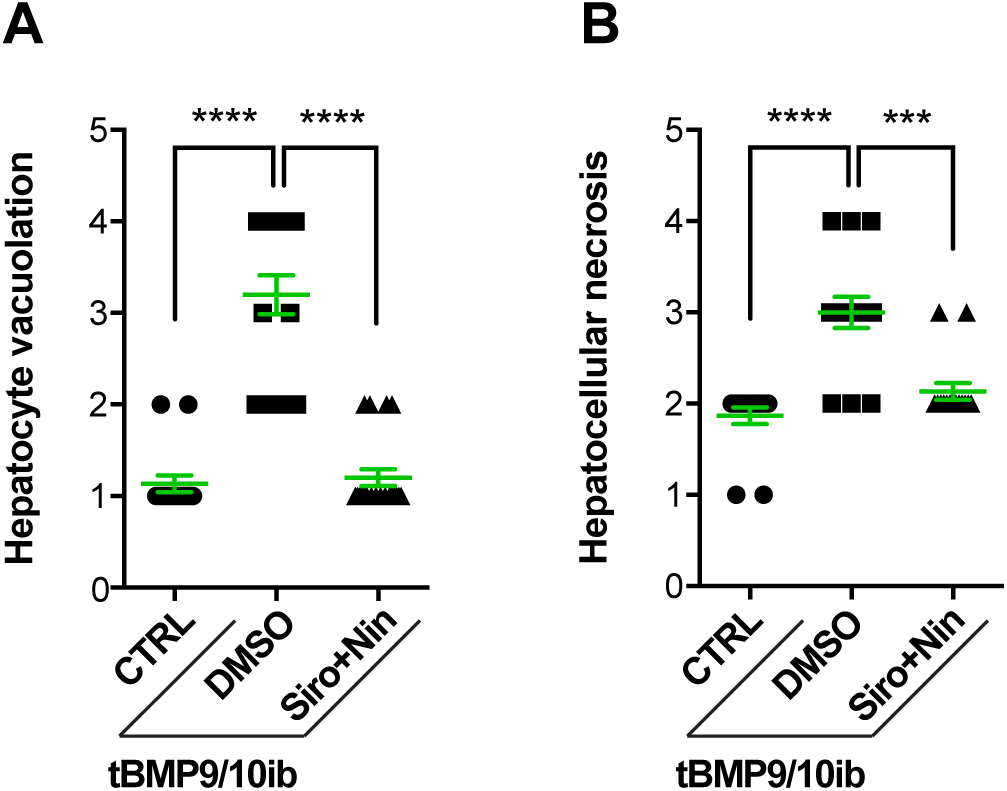
Siro+Nin attenuates hepatocyte toxicity in the liver of the tBMP9/10ib mice. Scatter plots of hepatocyte vacuolation **(A)** and hepatocellular necrosis **(B)** from liver of P9 control (CTRL) and tBMP9/10ib mice treated with DMSO and Siro+Nin, as in Figure 2A. Liver histological evaluation was performed according to the standardized semi-quantitative scoring system previously described (1). Data represent the mean of means ± SEM (5 random images of the liver per mouse, n = 3, 4, 4 mice for the CTRL, DMSO, and Siro+Nin groups, respectively); Kluskal-Wallis test, Dunn’s multiple comparison test; \*\*\**P*<0.001,\*\*\*\**P*<0.0001.

**Figure S8.**
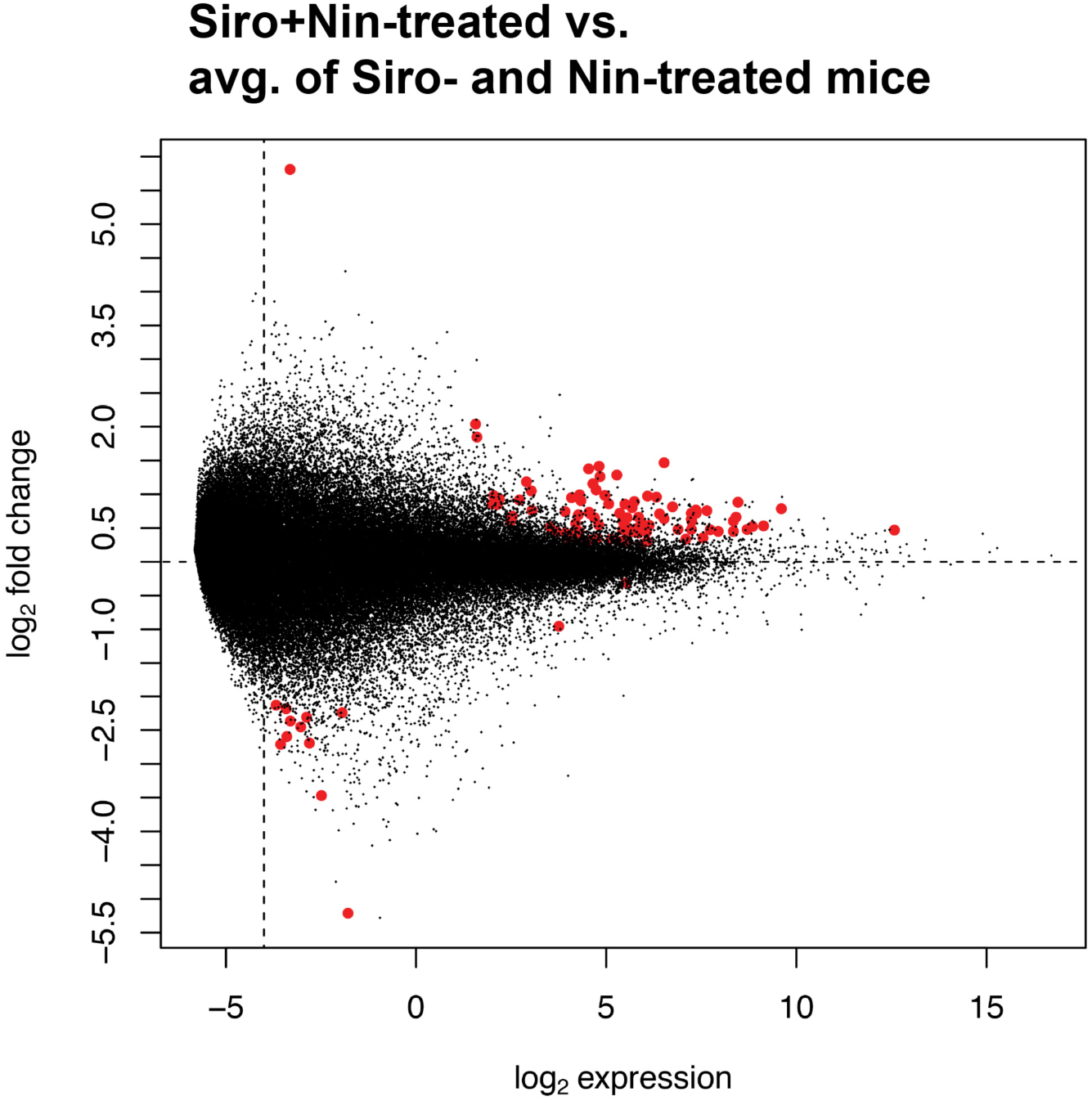
RNA-Seq quality control. We constructed an MA plot to visualize the effectiveness of the normalization and covariate control procedures employed during analysis of gene expression. The MA plot corresponds to the heatmap shown in Figure 6. The plot shows that the normalization and covariate adjustments employed were adequate for this dataset (most genes lie on a horizontal line corresponding to no change in gene expression with, as expected, more spread along the vertical for genes measured with low expression). Genes detected significant with a FDR≤1% are shown in red.

**Figure S9.**
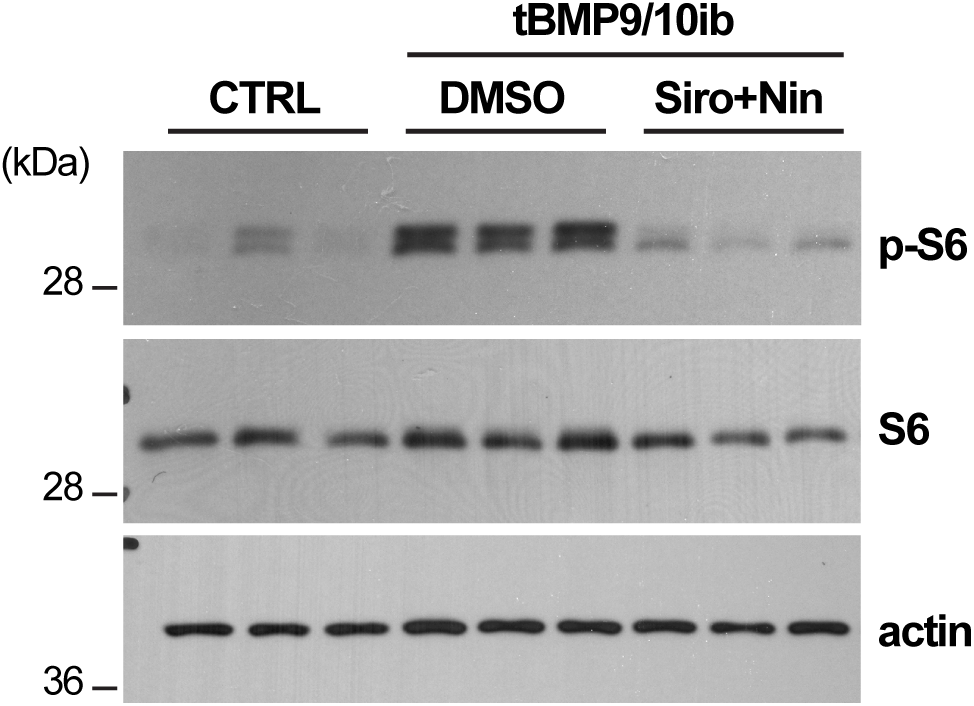
Siro+Nin inhibits S6 phosphorylation *in vivo* in tBMP9/10ib mice. P6 control (CTRL) and tBMP9/10ib mice were treated with DMSO or Siro+Nin, as in Figure 1A. Protein homogenates of whole liver were analyzed by WB using antibodies directed against the indicated proteins.

**Figure S10.**
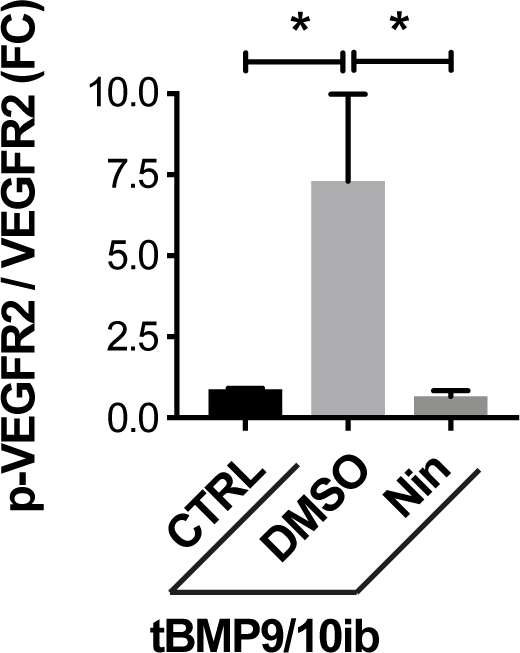
Nin prevents endothelial VEGFR2 overactivation *in vivo* in tBMP9/10ib mice. Densitometric analysis of the WB shown in Figure 8D quantifying phospho-VEGFR2 / total VEGFR2 ratio (p-VEGFR2/VEGFR2) in liver ECs isolated from CTRL and tBMP9/10ib mice treated with DMSO and Nin. FC, fold change. Data represent mean ± SEM (n = 5); one-way ANOVA, Tukey’s multiple comparison test; \**P*<0.05.

**Figure S11.**
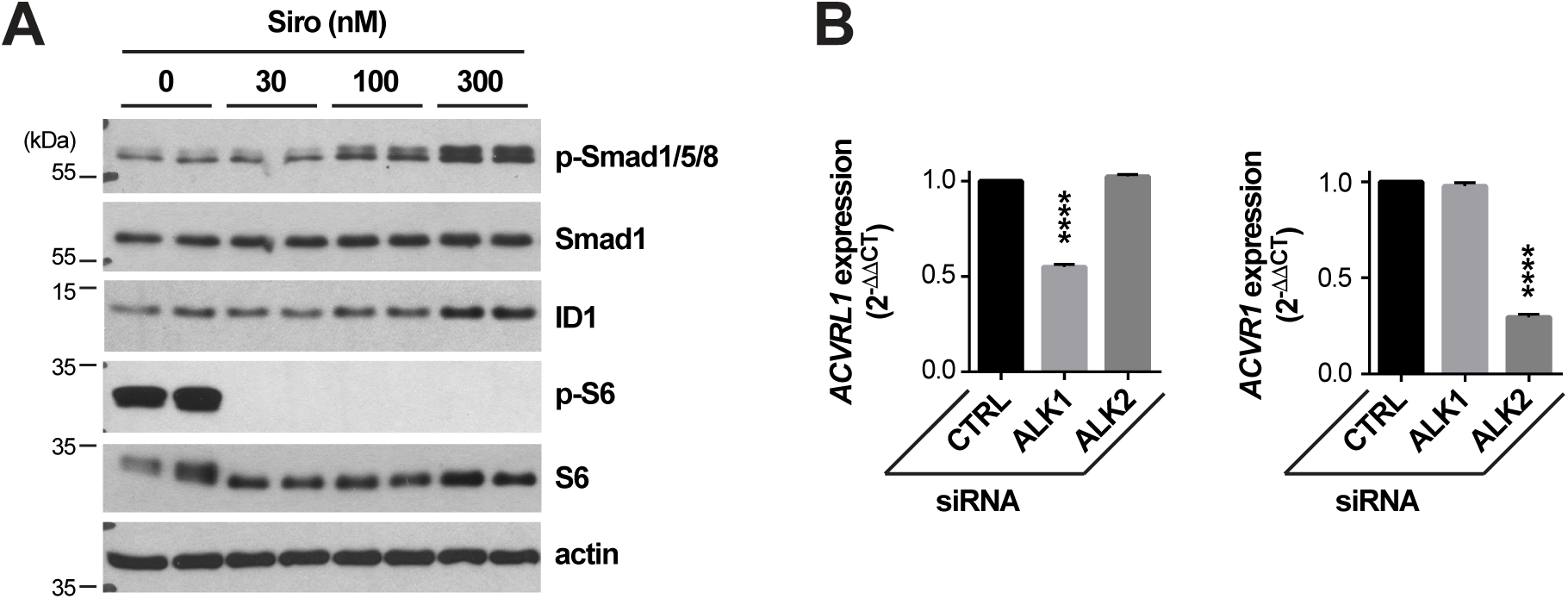
Siro rescues Smad1/5/8 signaling by activating ALK2. **(A)** C2C12 cells were treated for 3h with Siro in complete medium (conditioned for 2-3 days) and at the indicated concentrations. Cell extracts were analyzed by WB using antibodies directed against the indicated proteins. **(B)** Histogram showing *ACVRL1* (ALK1) and *ACVR1* (ALK2) expression (2^-^*^ΔΔ^*^Ct^) in HUVECs treated with CTRL or *ACVRL1*- and *ACVR1*-targeting siRNA. Data represent mean ± SEM (n = 3); one-way ANOVA, Tukey’s multiple comparison test; \*\*\*\**P*<0.0001.

**Figure S12.**
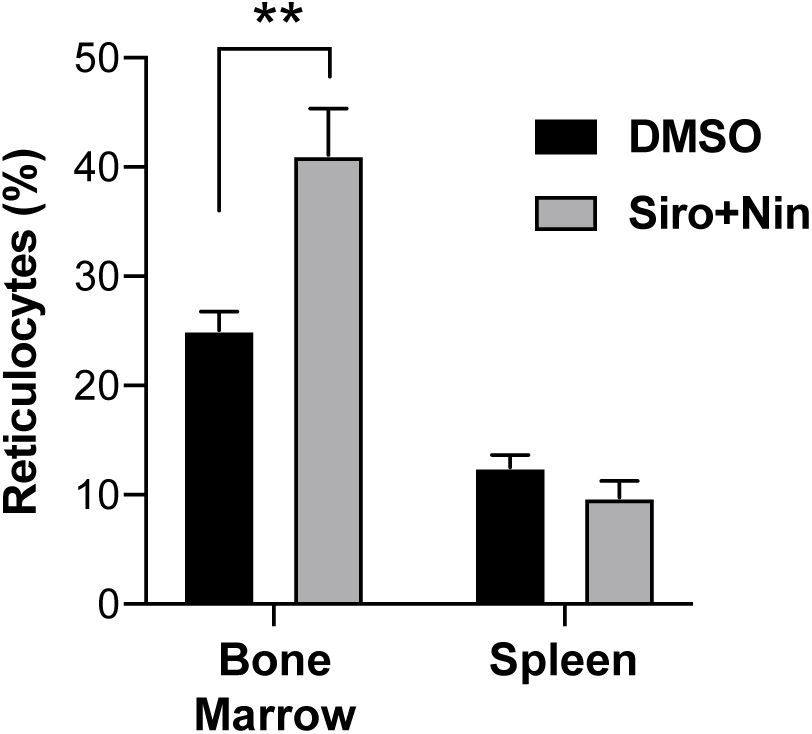
Siro+Nin increases the number of reticulocytes in the bone marrow of the tBMP9/10ib mice. Histogram shows reticulocyte counts determined in the bone marrow and spleen of P9 tBMP9/10ib mice treated as in Figure 2A with DMSO or Siro+Nin, and based on the expression levels of CD44 and Ter119, as described in (2). Data represent the mean of means ± SEM (n = 3 and 4 mice for the DMSO and Siro+Nin groups, respectively); two-way ANOVA, Bonferroni’s multiple comparison test; \*\**P*<0.01.

## SUPPLEMENTAL TABLE LEGENDS

**Table S1.** Transcripts differentially expressed upon synergistic drug treatment. Transcripts shown in the heatmap of Figure 6 are presented in table form. Selection was the same as the heatmap: FDR≤0.5%, log average expression >-5. The transcripts were annotated with Biomart and ensembl genes 94 annotations.

## SUPPLEMENTAL METHODS

### Reagents and antibodies

BMP9 (3209-BP) and VEGF_165_ (293-VE) were obtained R&D Systems. Sirolimus (Siro, S1039), nintedanib (Nin, S1010), and LDN-193189 (S2618) were from Selleck Chemicals. For WB, antibodies directed against p-S6 (4858), S6 (2217), p-Ser^2448^-mTOR (2971), mTOR (2983), p-VEGFR2 (2478), VEGFR2 (2479), p-Smad1/5/8 (13820), Smad1 (9743), p-ERK1/2 (9101), ERK1/2 (9102), p-p38 (4511), and p38 (9212) were obtained from Cell Signaling Technology. Anti-ID1 antibody was from BioCheck (BCH-1/195-14). Anti-actin antibody was from BD Transduction Laboratories (612656). For IHC, anti-PAI-1 (ab66705), anti-HIF-1α (ab1), and anti-Erg (ab92513) antibodies were obtained from Abcam, anti-Ter119 antibody from R&D Systems (MAB1125), and anti-p-S6 (4858), anti-p-VEGFR2 (2478) and p-Smad1/5/8 (13820) antibodies from Cell Signaling Technology. Alexa Fluor 594 anti-rabbit or anti-rat goat secondary antibodies were used for IHC (Molecular Probes).

### Transmammary-delivered immunoblocking of BMP9 and BMP10

For vascular disease induction in mouse pups, lactating dams were injected i.p. once at P3 with mouse monoclonal anti-BMP9 and anti-BMP10 antibodies (15 mg/kg bw, IgG2b, MAB3209; 15 mg/kg, IgG2a, MAB2926; R&D Systems, respectively), or with mouse monoclonal isotype controls (15 mg/kg, IgG2b, MAB004; 15 mg/kg, IgG2a, MAB003; R&D Systems) to obtain control mouse groups.

### Measurements of antibody levels by ELISA in neonatal mouse serum

ELISAs were performed as described before (3), with the following modifications. 96-well ELISA plates (Maxisorp, Nunc) were coated with 100 µL of 1 µg/mL recombinant BMP9 or BMP10 (R&D Systems) in coating buffer (15 mM K2HPO4, 25 mM KH2PO4, 0.1 M NaCl2, 0.1 mM EDTA, 7.5 mM NaN3) and incubated overnight at 4 °C. Plates were then washed 3 times with 0.05% Tween PBS (PBST) and blocked for 1 h at room temperature (RT) with 1% BSA in PBS. After washing 3 times with PBST, serial dilutions of individual mouse serum samples and reference mouse anti-BMP9 and anti-BMP10 IgGs (R&D Systems) (diluted in 1% BSA PBS) were prepared and 100 µL/well were incubated for 2 h at RT. After 3 more washes, 100 µL/well horseradish peroxidase (HRP)-conjugated goat anti-mouse secondary antibody (Southern Biotech, diluted 1:500 in 1% BSA PBS) was incubated for 1 h at room temperature. TMB substrate was added after 5 washes and the reaction was allowed to develop for 30 min at room temperature. The optical density was measured at 450 nm using a TECAN GENios Pro plate reader.

### Blue latex dye injection

P9 pups were processed for cardiac perfusion after euthanasia, following procedures described before (3). Briefly, the thorax and abdomen were opened. The left ventricles were injected manually with 600 µL of blue latex dye (BR80B, Connecticut Valley Biological Supply) using an insulin syringe U-100 (329652, BD Biosciences) and the right atrium was opened to drain the blood. After perfusion, liver, lungs, tongue, and palate were dissected, fixed in 4% paraformaldehyde (PFA), washed in phosphate-buffered saline (PBS) and either mounted in 80% glycerol (direct observation of the liver and lungs) or embedded in low gelling temperature agarose (A0701, Sigma) (tongue and palate). Blocks were dehydrated in methanol series and cleared with organic solvent (benzyl alcohol/benzyl benzoate, 1:1, Sigma). Images of the whole organ or tissue were acquired using an Olympus SZX7 stereomicroscope attached to an Olympus DP27 camera. Measurements and quantifications were performed using Fiji. For liver analysis, images from the same liver lobe and at the same location in the lobe were acquired at 1x magnification. The diameter of every hepatic vessel was measured at a distance of 2,000 µm from the margin of the lobe using the concentric circle tool plugin in Fiji. For the tongue and palate, images of the organs were acquired using 2x magnification and transversal lines were superimposed half-way in their anterior-posterior axis using the grid Fiji’s plugin, where the vessel diameter was measured. For the lungs, images of lobes were acquired and main vessels’ diameter was measured.

### IHC

To analyze the retinal vasculature, eyes were enucleated and fixed in 4% PFA for 20 min on ice and retinas were isolated and analyzed as before (3, 4). Briefly, retinas were dissected, cut four times to flatten them into a petal flower shapes, and fixed with methanol for 20 min on ice. After removing methanol, retinas were washed in PBS for 5 min on a shaker at room temperature, and blocked in blocking solution (0.3% Triton, 0.2% BSA in PBS) for 1 h on a shaker at room temperature. Retinas were then incubated in Alexa Fluor 488 isolectin B4 (I21411, Molecular Probes) and/or primary antibodies in blocking solution on a shaker overnight at 4°C. Retinas were then washed four times in 0.3% Triton in PBS for 10 min on a shaker, incubated with the corresponding secondary antibodies, washed four times in 0.3% Triton in PBS for 10 min followed by two washes in PBS for 5 min on a shaker and mounted in Vectashield (H-1000, Vector Laboratories).

Liver, heart, and spleen were fixed in 4% PFA for 24 h and sent for sectioning, H&E staining, and anti-HIF-1α IHC at HistoWiz. Unstained liver sections were dewaxed and antigen retrieval was performed. Sections (5 µm) were then washed in PBS for 5 min on a shaker at room temperature, and blocked in blocking solution (0.3% Triton, 0.2% BSA in PBS) for 1 h in a humid chamber at room temperature. Immunostaining was performed using primary antibodies in blocking solution in a humid chamber overnight at 4°C. Sections were then washed four times in 0.3% Triton in PBS for 10 min, incubated with the corresponding secondary antibody, washed four times in 0.3% Triton in PBS for 10 min followed by one incubation with DAPI (KPL 710300) in PBS and two washes in PBS for 5 min on a shaker before being mounted in Vectashield.

High resolution images for vascular network analysis were acquired using a Zeiss 880 laser confocal microscope. In cases where low magnification was enough for the analyses, an automated EVOS FL Auto 2 Invitrogen microscope was used. Measurements, quantifications, and 3D analysis were performed using Fiji. Briefly, low magnification images were acquired using a 4x and 10x lens to measure the retina vascular length (percentage of vasculature/retina length). A 20x lens was used to acquire images at the optic nerve to quantify AVM number, and AVM and vein diameter, and at the vascular plexus (3-5 fields per retina, between an artery and a vein) to quantify the vascular density. All measurements and qualifications were done using Fiji. The vascular density quantification was done by using the measure particles tool, working with 8-bit images, adjusting the threshold, and measuring the area occupied by the vasculature in a region of interest of 200 x 200 µm^2^. The bleeding area quantification was performed on 3 by 3 tiled image stacks using a 10x lens, and the 3D analysis by using the 3D project and the orthogonal view tools. Quantification of p-S6, p-VEGFR2, and p-Smad1/5/8 was performed by acquiring higher magnifications images using a 63x lens, analyzing 2 to 5 fields per retina and calculating the percentage of retinal vascular area occupied by immunostaining.

### CBC

Blood samples from tBMP9/10ib mice were obtained through heart puncture using a 0.3 mL insulin syringe (324910, BD Biosciences). Blood was transferred to a tube containing 5 mM EDTA (Invitrogen) and HCT, RBC number, and Hb levels were determined on a Beckman Coulter AcT analyzer, no more than 30 min after blood collection.

### Drug efficacy testing in adult Alk1 iKO mice

Tamoxifen was injected i.p. at 0.1 mg/g of body weight (b.w.) to 2-4 month-old R26^+/+^;*Alk1*^2f/2f^ (control group) and R26^CreER/+^;*Alk1*^2f/2f^ males and females on Day 0. Siro and Nin stock solutions were made in DMSO at 20 mg/mL and 10 mg/mL, respectively. The drugs were diluted in saline to 1X and adequate volume of drugs (Sir:Nin = 0.5:0.3 mg/kg bw) were injected i.p. once a day starting on Day 0. Saline containing 1% DMSO was used as vehicle control. Hb concentration in a drop of blood from tail snip was measured on Day 0, Day 7, and the last day (Day 9), using a Hb photometer (Hemopoint H2, STANBIO Laboratory). Any mice whose Hb levels were below 4 were subjected to termination on the day of measurement. At termination, mice were anesthetized by ketamine/xylazine (100/10 mg/kg bw), and GI tract was visually inspected and photographed. GI bleeding index was applied as 3 = severe; 2 = moderate; 1 = weak; 0 = none. Blood collected from abdominal vein was used for CBC on a HT5 Veterinary Hematology Analyzer (Heska). The whole body was fixed in formalin and bleeding was further inspected after collecting guts from stomach to rectum.

### Cell cultures and RNAi

HUVECs were isolated from umbilical veins obtained from anonymous donors and subcultured in 5% fetal bovine serum (FBS)-containing EC growth medium (ScienCell), as described before (5). C2C12 cells were obtained from ATCC and were maintained in Dulbecco’s Modified Eagle’s Medium (DMEM) supplemented with 10% FBS, penicillin, and streptomycin. RNA interference (RNAi) was performed in HUVECs using Accell SMARTpool siRNAs targeting *ACVRL1* (E-005302-00) and *ACVR1* (E-004924-00), or using an Accell non-targeting pool control (D-001910-10), and by following the manufacturer’s recommended protocols (Dharmacon).

### Liver EC isolation

ECs were isolated from P9 mouse livers using anti-CD31 microbeads (Miltenyi Biotec GmbH), following the manufacturer’s instructions.

### WB analyses and proteomic array

For WBs, protein extracts were processed as before (4), with the following modifications. Cultured cells, isolated liver ECs, or whole liver tissue were solubilized in RIPA buffer (20-188, EMD Millipore) supplemented with 1× Complete protease inhibitor mixture (11697498001, Roche). 5-20 µg of proteins (depending on the primary antibody used) were separated by SDS-PAGE, transferred onto nitrocellulose membranes, which were probed with primary and secondary antibodies. A standard ECL detection procedure was used. For proteomic array, whole liver tissue homogenates were prepared and protein extracts were applied to the arrays following the manufacturer’s instructions (ARY015, Proteome Profiler Mouse Angiogenesis Array Kit, R&D Systems).

### RT-qPCR

Three days after siRNA treatment, cells were harvested and RT-qPCR was performed on pellets of ∼100,000 cells per sample by using high capacity cDNA reverse transcription kit (Applied Biosystems) and by following the manufacturer’s protocol. PCR was performed using TaqMan assays targeting *ACVRL1* (Hs00953798_m1), *ACVR1* (Hs00153836_m1) and *GAPDH* (Hs03929097_g1) on ABI 7900 HT (Applied Biosystems, Life Technologies). *ACVRL1* and *ACVR1* expression levels were normalized to the reference gene *GAPDH*. Relative changes in gene expression were determined by the ΔΔCt method and using control values normalized to 1.0.

### RNA-Seq

Livers were rinsed with PBS and processed for RNA extraction using RNeasy mini kit (Qiagen), according to the manufacturer’s instructions. Total RNA quality was verified using Thermo Scientific NanoDrop and Agilent Bioanalyzer. RNA was processed for RNA-Seq at the Genomics Resources Core Facility, Weill Cornell Medical College, New York, NY. Briefly, cDNA conversion and library preparation were performed using the TrueSeq v2 Illumina library preparation kit, following manufacturers’ recommended protocols. Libraries were multiplexed 9 per lane and sequenced on 8 lanes of an Illumina HiSeq 4000 instrument. While the original and agreed upon design called for 8 lanes of sequencing in one run, the core facility sequenced these lanes in three separate runs, sequencing a subset of lanes twice, but not others. Analysis took this deviation into account by encoding which libraries had been sequenced with the altered design in the covariate matrix used for differential expression (the factor SEQUENCING_MOD encodes this information).

### Gene expression differential expression analysis

RNA-Seq transcript counts were analyzed with analysis scripts written with the MetaR languages (6). Briefly, gene expression analysis used Limma Voom to estimate statistics of differential expression while correcting for covariates. Depending on the analysis, covariates included drug treatment, antibody used to induce the disease phenotype, and sequencing modality (e.g., libraries sequenced in a single lane by the core facility, or sequenced on two lanes in separate runs, then pooled). Disease phenotype consists of HEALTHY (or CTRL) and DISEASE (or tBMP9/10ib).

Model used for Figure 6 (statistical analysis and heatmap):

We fit the RNASeq read counts with Limma Voom using the following model:

Count ∼ DRUG + ANTIBODY + SEQUENCING_MOD

Where Count is transcript read count; DRUG is COMBO=the Siro+Nin combination, SIRO=Siro alone, or NIN=Nin administered alone; ANTIBODY is tBMP9/10ib or CTRL, and SEQUENCING_MOD is sequencing modality. To identify genes whose expression indicate an interaction of the drug (synergistic effect), we selected genes for which the quantity (contrast for the statistical test of significance): COMBO - (SIRO+NIN)/ 2 significantly differed from zero. A contrast that differs from zero identifies genes whose expression changes are not additive with respect to the effect of each individual drug (because genes with additive drug effects would result in COMBO measuring the average expression across the drugs each administered alone). The MA plot for this differential expression analysis is shown in Figure S8.

### Human BOECs isolation and characterization

BOECs were isolated as described before (4) from an HHT2 patient carrying the T372fsX truncation [*ACVRL1* c.1112dup, first reported in Ref. (7)].

### Preparation of the standard and quality control solutions for liquid chromatography-mass spectrometry (LC-MS)

Standard stock solution of Siro and Nin were prepared in DMSO and diluted with ion-free water to prepare working solutions at concentrations of 0.01, 0.1, 1, and 10 µg/mL. Standard solutions in a concentration range between 0 and 1000 ng/mL for Siro and between 0 and 2500 ng/mL for Nin were prepared diluting the working solutions with either 50% MeOH in 5 mM ammonium formate water or 50% AcCN in 0.1% formic acid water, respectively. Quality control solutions used in the validation were prepared in the same manner as the standard solutions by using blank plasma collected from mice. All solutions were stored at −20°C before use.

### Sample preparation and LC-MS analysis

Samples (calibrator, control, and plasma) were mixed either with a 70:30 mix of acetonitrile and 0.1 M zinc sulfate in water or with acetonitrile (for Siro and Nin samples, respectively), followed by vortex mixing for 30 sec. After mixing, samples were left at room temperature for 10 min and centrifuged (10,000 x g) for 10 min. The supernatant was dried under nitrogen and the residue was reconstituted either with 50 µL of 5 mM ammonium formate/MeOH (50:50%, v/v) or 40 µL of water/acetonitrile, with formic acid (0.1% v/v). Reconstituted residue was vortexed for 10 sec and centrifuged at 13,000 x g for 10 min. The final extract was transferred into polypropylene vials, and then 10 µL was injected and analyzed on a LTQ XL™ Linear Ion Trap MS coupled to a vanquish UHPLC system (Thermo Scientific), with a Accucore™ Vanquish™ C18+ UHPLC Column (1.5 µm, 2.1 x 50 mm) at 45°C. A linear gradient of 50-100% methanol in 5 mM ammonium formate water (0.1% formic acid) or 2-95% acetonitrile was used for 6 and 10 min with flow rates of 0.2 and 0.4 mL/min, respectively. Data was analyzed using Xcalibur 3.1 (Thermo Scientific, USA), and Siro and Nin were identified through accurate mass measurements by comparison with pure standards. Peak areas of Siro and Nin were integrated and quantitated through the use of a calibration curve.

### Cardiac ultrasound measurements and analysis

During ultrasound examination, mice were lightly anesthetized using isoflurane (3% induction dose, 1.5% maintenance dose) plus oxygen (1.5 L/min). Body temperature was maintained in a physiological range using build-in heat pad. Chest and upper abdominal hair was removed by a chemical hair remover (Nair, Church & Dwight Co., Inc, NJ); then, ultrasound gel was applied to the skin to facilitate sound transmission and to reduce contact artifacts. Echocardiography was done with a Vevo 3100 instrument (VisualSonics, Canada). MX550D high-resolution transducer was used under mouse (small) cardiology mode (frequency: 40 MHz; acquisition frame rate: 232 lps; gain: 29 dB; depth of penetration 10 mm; width: 12.08 mm) and color doppler mode (frequency: 32 MHz; PRF: 30 KHz; acquisition frame rate: 33 lps; gain: 30 dB). Measurement was performed using build-in cardiac package of RV and PV function to obtain parameters of heart rate (beats per min, BPM), cardiac output (mL/min) by LV area long under B-mode and LV area under M-mode, and PAT (ms), PET (ms), PAT/PET ratio under PW Doppler mode (8).

